# DeepComBat: A Statistically Motivated, Hyperparameter-Robust, Deep Learning Approach to Harmonization of Neuroimaging Data

**DOI:** 10.1101/2023.04.24.537396

**Authors:** Fengling Hu, Alfredo Lucas, Andrew A. Chen, Kyle Coleman, Hannah Horng, Raymond W.S. Ng, Nicholas J. Tustison, Kathryn A. Davis, Haochang Shou, Mingyao Li, Russell T. Shinohara, The Alzheimer’s Disease Neuroimaging Initiative

## Abstract

Neuroimaging data from multiple batches (i.e. acquisition sites, scanner manufacturer, datasets, etc.) are increasingly necessary to gain new insights into the human brain. However, multi-batch data, as well as extracted radiomic features, exhibit pronounced technical artifacts across batches. These batch effects introduce confounding into the data and can obscure biological effects of interest, decreasing the generalizability and reproducibility of findings. This is especially true when multi-batch data is used alongside complex downstream analysis models, such as machine learning methods. Image harmonization methods seeking to remove these batch effects are important for mitigating these issues; however, significant multivariate batch effects remain in the data following harmonization by current state-of-the-art statistical and deep learning methods. We present DeepCombat, a deep learning harmonization method based on a conditional variational autoencoder architecture and the ComBat harmonization model. DeepCombat learns and removes subject-level batch effects by accounting for the multivariate relationships between features. Additionally, DeepComBat relaxes a number of strong assumptions commonly made by previous deep learning harmonization methods and is empirically robust across a wide range of hyperparameter choices. We apply this method to neuroimaging data from a large cognitive-aging cohort and find that DeepCombat outperforms existing methods, as assessed by a battery of machine learning methods, in removing scanner effects from cortical thickness measurements while preserving biological heterogeneity. Additionally, DeepComBat provides a new perspective for statistically-motivated deep learning harmonization methods.

## 1 Introduction

There is increasing need for larger sample sizes in human magnetic resonance imaging (MRI) studies to detect small effect sizes, train accurate prediction models, improve generalizability, and more. This has led to more interest in multi-batch studies, where subjects are imaged across multiple sites or scanners and then aggregated together (Bethlehem et al., 2022; Casey et al., 2018; Di Martino et al., 2014; Marek et al., 2022; Mueller et al., 2005; Trivedi et al., 2016; Van Essen et al., 2013). multi-batch studies overcome limitations of single site studies, which are often unable to recruit sufficiently large or representative samples to achieve study goals; however, multi-batch study designs introduce non-biological, technical variability between subjects imaged from different batches due to differences in acquisition, scanner manufacturer, magnet strength, post-processing, and more (Badhwar et al., 2020; Han et al., 2006; Jovicich et al., 2006; Takao et al., 2014, 2011). Such technical variability is often referred to as “scanner effects” or “batch effects” and, if not appropriately addressed, may result in invalid, non-reproducible, or non-generalizable study results. Post-acquisition removal of these batch effects, known as image harmonization, is a promising approach for mitigating these issues (Hu et al., 2023).

Harmonization of image-derived features, such as cortical thicknesses, functional connectivity values, radiomics features, and more has been extensively studied. Fortin et al. (2017) showed that the ComBat model, adapted from the genomics setting, could effectively remove batch effects by modeling them univariately as additive differences in means and as multiplicative differences in variances of residuals (Johnson et al., 2007). This model has also been extended to unique data settings, such as those where covariate effects are non-linear, longitudinal data is present, decentralized learning is required, multiple batch variables should be corrected for, or traveling subjects are available (Bayer et al., 2022; Bostami et al., 2022; Chen et al., 2022b; Horng et al., 2022; Maikusa et al., 2021; Pomponio et al., 2020). In applied studies, ComBat-family methods have been widely used and shown to improve inference and generalizability of results (Acquitter et al., 2022; Bartlett et al., 2018; Bourbonne et al., 2021; Crombé et al., 2020; Fortin et al., 2018; Marek et al., 2019; Yu et al., 2018). This may be especially true in mass univariate inference settings, where biological effects are modeled at the individual feature level, since this setting matches the data assumptions made by the ComBat model.

However, in studies where feature-level data is used in a highly multivariate manner, univariate harmonization approaches may be insufficient. For example, as imaging researchers have become more interested in complex prediction efforts, multivariate feature datasets are used as inputs to predict an outcome of interest. In these settings, state-of-the-art machine learning (ML) algorithms are often used as powerful approaches that are able to jointly leverage the multivariate distribution of features, accounting for complex non-linear and interaction effects (Hu et al., 2023; Koutsouleris et al., 2014; Smith et al., 2017; Wager et al., 2013). Batch effects that exist in the interactions between features may also be picked up by these ML algorithms, which can lead to decreased generalizability of these models and overfitting of model parameters on batch effects, especially when batch status is a relevant confounder for the outcome. Thus, recent efforts in feature-level harmonization have attempted to detect and mitigate such multivariate batch effects.

From the statistical perspective, recently proposed methods for multivariate harmonization have included CovBat (Chen et al., 2022a), Bayesian factor regression (BFR, Avalos-Pacheco et al., 2022), and UNIFAC (Zhang et al., 2022). Like ComBat, these models assume batch effects can be effectively modeled through the combination of low-rank additive and multiplicative effects. However, instead of modeling batch effects solely in a univariate manner, CovBat additionally assumes batch effects to be present in the covariance structure of model residuals, while BFR and UNIFAC assume additive batch effects to be present in the direction of multivariate latent factors. Additionally, while ComBat, CovBat, and UNIFAC all seek to ultimately produce a dataset of harmonized features, BFR instead learns a low-dimensional representation of the original features where batch effects have been removed; BFR does not map this low-dimensional representation back to the feature space.

From the deep learning perspective, feature-level multivariate harmonization methods have leveraged the conditional variational autoencoder (CVAE) architecture, an adaptation of the standard variational autoencoder that attempts to disentangle the latent space distribution from covariates of interest (Kingma and Welling, 2014; Sohn et al., 2015). These models include diffusion CVAE (dcVAE, Moyer et al., 2020) and goal-specific CVAE (gcVAE, An et al., 2022). In dcVAE, an encoder is used to embed vector representations of diffusion MRI data as latent space distributions, and the encoder is penalized when batch-specific information is present in the latent space representation. Then, the decoder is given these latent space distributions along with explicit batch information and trained to reconstruct the original input. Through this process, dcVAE assumes that the encoder can learn to remove batch effects and the decoder can accurately reconstruct the original data, but with batch effects removed. However, An et al. (2022) noted that dcVAE may inadvertently remove biological information of interest. They proposed gcVAE could recover this biological information by fine-tuning the dcVAE decoder such that the decoder could not only accurately reconstruct the input but could also retain biological information of interest in the reconstruction. gcVAE encourages this behavior by adding an additional pre-trained neural network classifier to the end of dcVAE that attempts to use decoder output to predict biological covariates of interest. Classifier success is rewarded in the loss function.

Finally, there have been extensive efforts in performing image-level harmonization, where batch effects are removed from raw MRI images instead of from image-derived features (Bashyam et al., 2022; Cackowski et al., 2021; Fatania et al., 2022; Fetty et al., 2020; Hiasa et al., 2018; Karras et al., 2019; Liu et al., 2021; Modanwal et al., 2020; Tian et al., 2022; Yao et al., 2022; Zhao et al., 2019; Zhu et al., 2017; Zuo et al., 2021). These methods attempt to disentangle biological variability from technical variability through the use of generative adversarial networks (GANs) or convolutional autoencoder-style models similar to the CVAE networks describe above.

In methods based on cycle-consistency GAN (CycleGAN), two generator-discriminator pairs are trained together (Hiasa et al., 2018; Modanwal et al., 2020; Zhao et al., 2019; Zhu et al., 2017). The first pair seeks to make images from the first batch look like those from the second batch, while the second pair seeks to make images from the second batch look like those from the first batch. Importantly, a cycle-consistency constraint is enforced, such that when images from the first batch are cycled to the second batch and then back to the first batch, these cycled images are consistent with the original raw images. The same constraint is enforced for images from the second batch.

For autoencoder-based methods applied at the image level, ideas similar to CVAE are used, where methods seek to decompose images into batch-invariant content representations in the encoding step, and then in the decoding step, inject these content representations with batch information necessary for reconstruction (Bashyam et al., 2022; Cackowski et al., 2021; Fatania et al., 2022; Fetty et al., 2020; Karras et al., 2019; Liu et al., 2021; Tian et al., 2022; Yao et al., 2022; Zuo et al., 2021). Broadly, in this class of harmonization methods, batch discriminators can be used to impose penalties on embedding batch information into the latent space, GANs can be used as decoder modules in order to achieve more realistic reconstructions, and cycle-consistency losses can be imposed to encourage disentanglement of batch and content. Importantly, instead of in feature-level CVAE, where batch information is merely concatenated to the latent space representations at the decoding stage, in image-level autoencoder-based methods, adaptive instance normalization (AdaIN) is commonly used to inject batch information (Huang and Belongie, 2017). AdaIN showed that in convolutional autoencoders, where latent space representations consist of convolutional feature maps where each feature map can be thought to indicate the locations and strength of that feature in the input image, arbitrary style transfer can be performed in the latent space by shifting and rescaling each feature map such that its mean and variance match those of the corresponding feature map in the desired style. Intuitively, AdaIN proposes that style, or batch status, is largely encoded in the first two moments of the latent space representation and that globally changing these moments within each feature map results in style transfer. Non-convolutional autoencoders have been shown to similarly encode style information in the latent space, at least with respect to the first moment, in single-cell RNA sequencing (Lotfollahi et al., 2019).

Notably, deep learning harmonization methods designed for both feature-level and image-level data make a number of strong implicit assumptions. Firstly, deep learning harmonization methods tend to directly use model outputs from the harmonization step as the resulting harmonized data – unmodeled residual terms are unaccounted for, as well as any batch or biological effects that may exist in these residuals. Implicitly, this makes the strong assumption that the deep learning method is able to achieve perfect or nearly-perfect model fit – that is, the reconstruction loss and cycle-consistency loss for autoencoders and cycle-consistency GANs, respectively, is zero or nearly zero. This is in contrast to statistical harmonization methods, which tend to estimate batch effects within unmodeled residual terms as a difference in scale; the residuals are rescaled and added back to the model-based biological effects to produce the resulting harmonized data. Secondly, deep learning harmonization methods assume that batch and biological effects can be completely disentangled through loss function optimization and choice of network architecture. While this may be easily achievable in isolation, complete disentanglement may be challenging to achieve in conjunction with the implicit nearly-perfect model fit assumption. Finally, deep learning harmonization methods do not explicitly take into account that biological covariates may be imbalanced across batches – in such cases, some population-level differences across batches may actually be due to true biological differences and therefore should not be removed.

In this manuscript, we propose a novel deep learning harmonization method, called DeepComBat, that is designed to effectively remove multivariate batch effects in a statistically-informed manner. Compared to statistical methods such as ComBat and CovBat, DeepComBat promises removal of complex, non-linear, and multivariate batch effects from the raw data in a way that mitigates detection of batch effects using highly multivariate methods. Compared to other deep learning methods, DeepComBat avoids making the assumptions described above – unmodeled residual terms are explicitly accounted for and corrected, a completely disentangled latent space is not required, and model-based batch effects are removed conditional on biological covariates that may be confounders. To the best of our knowledge, DeepComBat is the first deep learning harmonization method that explicitly accounts for confounders or unmodeled residuals. Additionally, DeepComBat hyperparameters can be tuned manually based on readily-accessible latent space summary statistics, and DeepComBat can be thought to have a form of “double-robustness” such that even with poor model fit, reasonable harmonization can still be achieved.

We apply DeepComBat to cortical thickness measurements acquired by the Alzheimer’s Disease Neuroimaging Initiative (ADNI) and compare our results to those of other feature-level harmonization methods where open-source code was available, namely: ComBat, CovBat, dcVAE (modified for non-diffusion setting), and gcVAE. We find that, compared to other methods, DeepComBat-harmonized data retains biological information of interest while containing minimal batch information as assessed by a number of ML methods. Our results demonstrate the advantage of incorporating statistical ideas into deep learning methods in order to more effectively perform multivariate harmonization.

## 2 Methods

### 2.1 ADNI dataset and preprocessing

We included 663 unique subjects (381 males) from the Alzheimer’s Disease Neuroimaging Initiative (ADNI, http://adni.loni.usc.edu/). For each subject, the most recent T1-weighted (T1w) imaging acquired during the ADNI-1 phase was used; all included images were acquired between July 2006 and August 2010. Informed consent was obtained for all subjects in the ADNI study. Institutional review boards approved the study at all of the contributing institutions.

For the purposes of this study, we define two batches based on scanner manufacturer – the first batch consists of all subjects imaged on scanners manufactured by Siemens Healthineers (n = 280) and the second batch consists of all subjects imaged on scanners manufactured by either Philips Medical Systems (n = 96) or GE Healthcare (n = 287). These two batches were chosen based on findings by Fortin et al. (2018), who showed marked cortical thickness differences were present between images from Siemens and non-Siemens scanners, while minimal differences were present between images from Philips and GE scanners. Philips and GE manufacturers were combined into one batch to allow for improved estimation of the batch effects between Siemens and non-Siemens scanners.

Additionally, we define age, sex, and Alzheimer disease status (cognitively normal, CN; late mild cognitive impairment, LMCI; Alzheimer disease, AD) as biological covariates of interest that may confound the relationship between batch status and T1w imaging – these covariates are known to affect brain structure and also may be associated with scanner manufacturer through differing population demographics across sites. Subject demographics at time of most recent acquisition are presented in Table 1, stratified by these two batches. Notably, there are marked differences in the distribution of sex across the two batches, suggesting that confounding of batch status by subject demographics is plausible, and estimation of batch effects should be conditioned on subject demographics.

**Table 1.** Patient demographics at time of acquisition, stratified by batch.

| Characteristic | Philips/GE, N = 383<sup>1</sup> | Siemens, N = 280<sup>1</sup> |
| --- | --- | --- |
| Age | 77.0 (6.9) | 77.8 (6.6) |
| Sex |  |  |
| Male | 234 (61%) | 147 (52%) |
| Female | 149 (39%) | 133 (48%) |
| Diagnosis |  |  |
| Cognitively Normal | 115 (30%) | 82 (29%) |
| Late Mild Cognitive Impairment | 185 (48%) | 139 (50%) |
| Alzheimer Disease | 83 (22%) | 59 (21%) |
| Mini-Mental State Examination Score | 25.2 (4.9) | 24.8 (5.5) |
| <sup>1</sup> Mean (SD); n (%) |  |  |

Processing of these data was carried out using the Advanced Normalization Tools (ANTs) longitudinal single-subject template pipeline (Tustison et al., 2019). Briefly, we first downloaded raw T1w images from the ADNI-1 database, which were acquired using MPRAGE for Siemens and Philips scanners and using a works-in-progress version of MPRAGE for GE scanners (Jack Jr. et al., 2010). For each subject, we estimated a single-subject template using all image timepoints, and applied rigid spatial normalization to this template for each timepoint image. Then, each normalized timepoint image is processed using the single-image cortical thickness pipeline consisting of 1) brain extraction (Avants et al., 2010), 2) denoising (Manjón et al., 2010), 3) N4 bias correction (Tustison et al., 2010), 4) Atropos *n*-tissue segmentation (Avants et al., 2011), 5) and registration-based cortical thickness estimation (Das et al., 2009). Finally, for our analyses, we used cortical thickness values for the 62 Desikan-Killiany-Tourville atlas regions such that the feature matrix we sought to harmonize was of dimension 663 × 62 (Klein and Tourville, 2012). Scan metadata were determined based on information contained within the Digital Imaging and Communications in Medicine (DICOM) headers for each scan.

### 2.2 ComBat model

We first review the ComBat (Combatting Batch Effects) model, which models additive and multiplicative batch effects in an empirical Bayes framework (Fortin et al., 2017; Johnson et al., 2007). This model is used as a building block for DeepComBat. For each subject, let ***y****_ij_* = [*y_ij_*_1_, …, *y_ijk_*, …, *y_ijp_*]^τ^ represent the *p* × 1 vector of feature-level information for that subject, where each *y_ijk_* is a scalar. In this notation, *i* = 1,2, …, *B* indexes batch; *j* = 1,2, …, *n_i_* indexes subjects within batch *i*, where *n_i_* is the number of subjects acquired in batch *i*; and *k* = 1,2, …, *p* indexes features, where *p* is the total number of features. First, ComBat is fit on each feature individually using the following model:

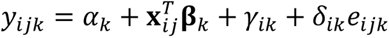

where *α_k_* is the vector of shared intercepts across batches; ***x****_ij_* is the vector of subject-specific biological covariates; **β***_k_* is the vector of regression coefficients for the covariates; *γ_ik_* is the vector of mean batch effects for batch *i* conditional on the covariates; and *δ_ik_* is the vector of multiplicative batch effects on the residuals. ComBat assumes the errors, *e_ijk_*, are distributed 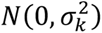.

For each individual feature, least-squares estimates 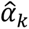 and 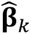 are obtained. Then, to estimate batch effects using empirical Bayes, ComBat assumes the additive batch effects, *γ_ik_*, are drawn from a normal distribution prior and the multiplicative batch effects, *δ_ik_*, are drawn from an inverse gamma distribution prior. Hyperparameters for these priors are estimated via method of moments using data across all features. Next, for each feature-level, empirical Bayes estimates, 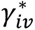 and 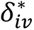, are obtained as the means of their corresponding posterior distributions. This results in shrinkage estimators for both the additive and multiplicative batch effects such that these effects can be well-estimated even when within-batch sample size is small. Finally, estimated batch effects are removed using the following equation:

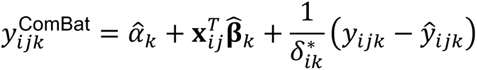

where 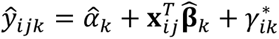 is the subject-specific mean as estimated by the ComBat model.

### 2.3 DeepComBat method

The DeepComBat method consists of three steps: 1) normalization, 2) CVAE training, and 3) harmonization. A broad overview of the method is given here, and further details are described in the following sections. First, the normalization step seeks to transform raw data such that the CVAE training step may converge more quickly. Then, the CVAE attempts to learn a latent space representation of the input data that contains rich subject-specific information, but contains fewer batch effects than the input data. In this step, the CVAE also learns to use this latent space representation along with explicit batch and biological information to reconstruct the data. Next, since these reconstructions are imperfect and batch effects may also be present in the reconstruction residuals, these residuals are harmonized using ComBat. Additionally, batch effects in the latent space are harmonized using ComBat, and the CVAE decoder uses this harmonized latent space along with the reference batch covariate to generate harmonized subject-specific means. Finally, the harmonized residuals are added to these harmonized means to obtain the final harmonized data. Overall, DeepComBat partitions batch effects into three components – the latent space, the CVAE decoder, and the reconstruction residuals. Each of these components is individually harmonized and then combined to produce the final DeepComBat-harmonized data. Notably, although DeepComBat effectiveness is demonstrated here between two batches, the code and architecture allow for harmonization between more than two batches without the need for alteration.

#### 2.3.1 Normalization

As in the above notation, let ***y****_ij_* = [*y_ij_*_1_, …, *y_ijk_*, …, *y_ijp_*]^τ^ represent subject *ij*’s cortical thickness vector, where *k* indexes features, and ***x****_ij_* represent the vector of subject-specific biological covariates. Additionally, let *b_ij_* represent that subject’s batch covariate.

In the normalization step, all biological covariates are linearly shifted and scaled across all *ij* subjects such that they range between 0 and 1. Batch covariates are indicators and are thus already in this range. Additionally, each feature is standardized across all *ij* subjects such that the overall mean for that feature is 0 and the variance is 1. CVAE training and harmonization steps use this normalized data; however, the linear transformations of features are stored such that they can be inverted, and the harmonized output will remain in the original feature space.

Normalization of biological covariates is theoretically unnecessary, but practically may allow for faster convergence of the DeepComBat CVAE since default deep learning weight initializations and hyperparameters are designed for inputs approximately in the range [0, 1]. Standardization of features, however, is necessary. The DeepComBat CVAE loss function, discussed further below, includes the mean-squared error (MSE) loss – if features are on drastically different scales, DeepComBat will prioritize reconstruction of features with large magnitudes at the expense of features with small magnitudes. Standardization allows for errors in reconstruction of all features to contribute to the loss function relatively equally. Additionally, standardization may provide practical benefits; as above, it may allow for faster convergence, and secondly, DeepComBat hyperparameters used in this study for standardized ADNI cortical thickness dataset may be more generalizable to other standardized datasets.

#### 2.3.2 Architecture

In the CVAE training step, normalized ADNI cortical thickness data are passed through a standard, fully-connected CVAE-style model with the architecture shown in Figure 1. For architectural hyperparameters, the latent space was empirically chosen to be approximately one-fourth the size of the input vector, rounded to the nearest power of 2 – in practice, latent spaces approximately one-eight or one-half the size of the input vector also performed similarly. Four hidden layers were used on either side of the latent space to allow for sufficient complexity of the encoder to learn meaningful latent space representations with minimal batch effects and of the decoder to incorporate batch effects in reconstruction. Hidden layer sizes were defined such that each size was approximately halfway between the size of the layers before and after.

**Figure 1:**
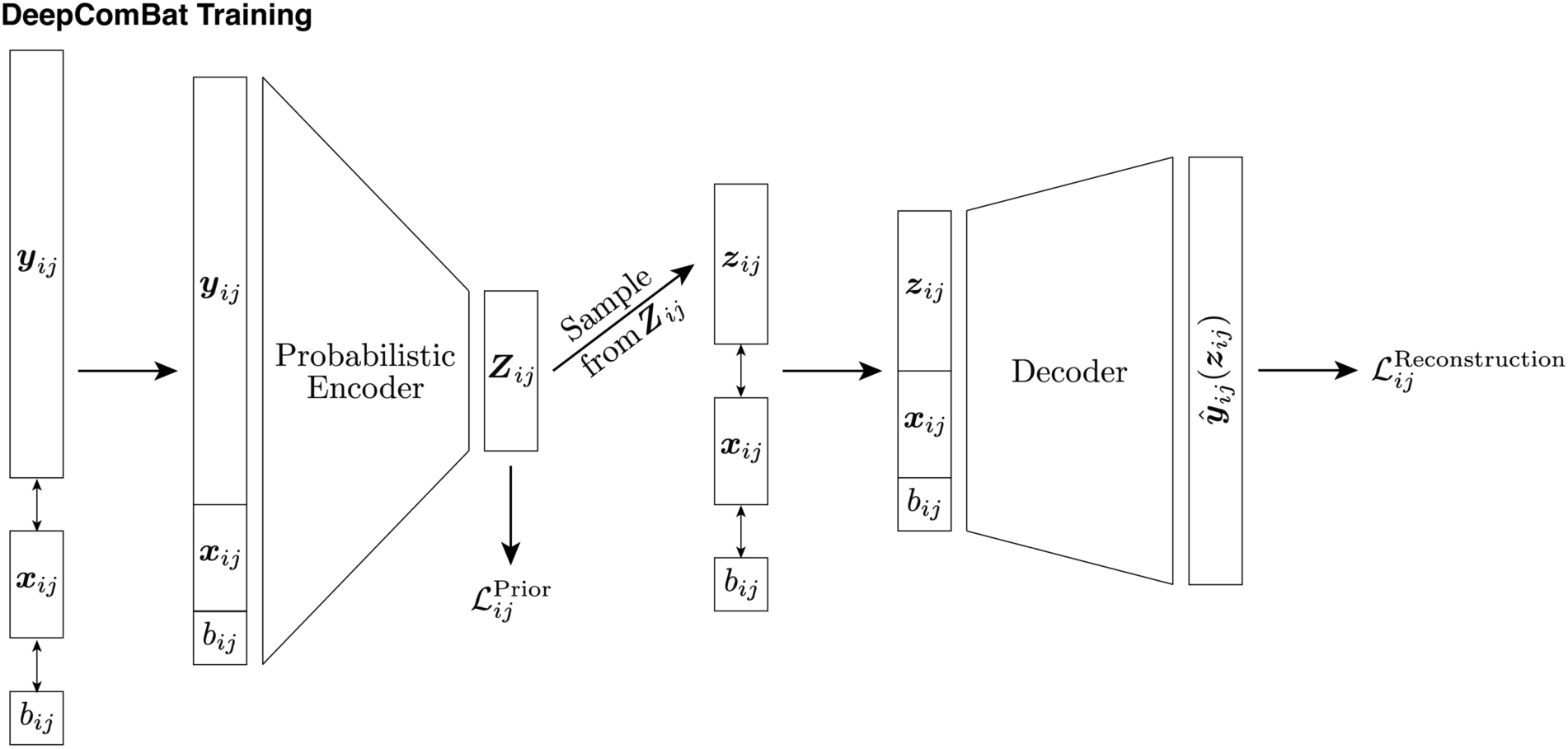
DeepComBat CVAE architecture and loss functions used during training. Notation corresponds to that in the main text.

One iteration through the CVAE for one subject is as follows. First, let the encoder input be defined as the column-wise concatenation of the column vectors ***y****_ij_*, ***x****_ij_*, and *b_ij_*. This encoder input is passed through successive hidden layers until it is eventually encoded into two 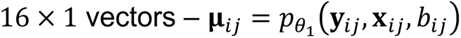 and 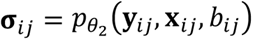, where 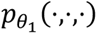 and 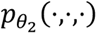 represent the encoder functions with neural network parameters *θ*_1_ and *θ*_2_, respectively. These vectors together define a multivariate normal random variable, **Z***_ij_* ~ *N* (**μ***_ij_*,diag(**σ***_ij_*)). This random variable is the output of the encoder and can be thought of as subject *ij*’s latent space representation. Next, to begin the decoding step, a sample is drawn from this random variable using the reparameterization trick in order to obtain **z***_ij_* (Kingma and Welling, 2014). As with the encoder input, this sample is column-wise concatenated with ***x****_ij_* and *b_ij_* to produce the decoder input. Then, it is passed through the decoder hidden and output layers to obtain a reconstructed feature vector, 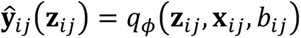, where *q_ϕ_*(·,·,·) is the decoder function with neural network parameters *ϕ*. Note that this reconstructed feature vector is a function of the sample from the random variable **Z***_ij_* and thus changes each time subject *ij* is passed through the CVAE.

Thus, the latent space distribution, **Z***_ij_*, is a function of the features, ***y****_ij_*, as well as on the covariates ***x****_ij_* and *b_ij_*. Similarly, the reconstructed feature vector, 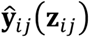 is a function of the latent distribution through **z***_ij_*, as well as on the covariates ***x****_ij_* and *b_ij_*. Additionally, by giving the decoder random samples from the latent space distribution, the decoder learns that probabilistically-nearby points in the latent space should be mapped to similar outputs in the feature space. That is, the decoder learns to reconstruct the features such that 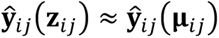. The risk of overfitting by the decoder is also minimized, as this random sampling functions as a form of data augmentation with respect to the decoder.

#### 2.3.3 Loss function

The loss function was defined to be the standard CVAE loss function which consists of an autoencoder reconstruction loss component and a Kullback-Leibler (KL) divergence loss component (Kingma and Welling, 2014; Sohn et al., 2015). In the DeepComBat CVAE, this loss function is implemented for each subject as follows:

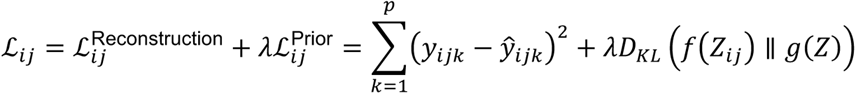

where 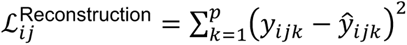 is the reconstruction component, 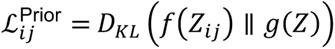 is the KL divergence component, and *λ* is a hyperparameter to weight the relative importance of the two components. The KL divergence component measures the difference between *f*(*Z_ij_*), which is the probability density function of the multivariate normal latent space distribution for subject *ij*, *N* (**μ***_ij_*,diag(**σ***_ij_*)), and *g*(*Z*), which is defined in DeepComBat to be the probability density function of the standard multivariate normal distribution, *N*(**0, I**). The overall loss function is defined as the sum over all subjects: 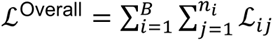.

The KL divergence term can be thought to enforce a standard normal Bayesian prior on the latent space, where *λ* represents the strength of the prior. For large *λ*, all latent space distributions converge to the uninformative prior, *N* (**μ***_ij_*,diag(**σ***_ij_*)) → *N*(**0, I**), and for *λ* close to 0, latent space distributions converge to the point estimate for the mean, *N* (**μ***_ij_*,diag(**σ***_ij_*)) → **μ***_ij_*. Thus, the KL divergence term allows for regularization of the latent space as well as encourages removal of information that is unnecessary for reconstruction from the latent space. In the DeepComBat CVAE, since biological and batch covariates are explicitly given to the decoder, optimal latent space representations should contain no information about these covariates and instead encode richer, subject-specific information. Practically, this complete independence may be unrealistic to achieve. Importantly, while biological and batch covariates are used as inputs for both the encoder and the decoder, the CVAE is not rewarded for including information about these covariates in the loss function. This design choice prevents the CVAE from introducing bias, but still allows the model to learn multivariate batch effects conditional on potential biological confounders.

#### 2.3.4 Optimization and hyperparameter tuning

This CVAE loss function is known to have the potential to suffer from KL vanishing, also referred to as posterior collapse, where a local minimum of the loss function is reached and the model cannot improve (Bowman et al., 2016). In KL vanishing, the encoder learns to collapse all latent space representations to the standard normal prior such that the KL component of the loss function is nearly zero, and the decoder is given total noise and is therefore unable to learn anything in order to make progress towards further minimizing the loss. To minimize risk of posterior collapse in the DeepComBat CVAE, we utilize a cyclic annealing optimization schedule (Fu et al., 2019). In this schedule, *λ* is gradually increased from 0 to the goal final KL divergence weight multiple times over the course of model training. This provides opportunities for the optimizer to escape local minimum when *λ* is small and allows for progressive learning of more meaningful latent representations across cycles.

In DeepComBat, we perform manual hyperparameter tuning, described in more detail in the Results section, to determine our desired final *λ*_Final_ = 0.1. The goal in tuning *λ*_Final_ is to impose a prior that is strong enough to regularize the latent space and Euclidean distance between latent space representations are meaningful, but weak enough to allow for rich, subject-specific information to be encoded in the latent space in order to produce high-quality reconstructions.

Note that, in contrast to similar CVAE-based harmonization methods like dcVAE, gcVAE, and a number of image-based methods which require a KL divergence component hyperparameter such that latent space distributions are independent of batch, the DeepComBat *λ*_Final_ is instead only used to regularize the latent space and reduce the amount of batch information in the latent space, if possible. However, substantial remaining batch information in the DeepComBat latent space is allowed, which enables easier hyperparameter tuning.

Using this *λ*_Final_, we first pre-train the CVAE for 5 epochs with *λ* = 0, then perform cyclic annealing over 30 epochs where one cycle is 5 epochs and *λ* increases linearly from 0 to *λ*_Final_ within each cycle, and finally train the CVAE for 5 epochs with the desired *λ* = *λ*_Final_. Optimization was performed using the Adam optimizer with learning rate of 0.01, chosen to increase the initial rate of model convergence (Kingma and Ba, 2017). Within epochs, data was passed to the CVAE in mini-batches of 64 subjects.

#### 2.3.5 Harmonization

Once the CVAE model has been trained, harmonization can be performed on the latent space, the CVAE decoder, and the reconstruction residuals, as shown in Figure 2. In the latent space, each subject’s noisy latent space distribution, **Z***_ij_*, is converted to the noiseless latent space mean vector, **μ***_ij_*. Then, across all *ij* subjects, the ComBat model described above is fitted using both batch and biological covariates to harmonize the latent space. Let each ComBat-harmonized latent space representation be denoted as: 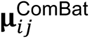.

**Figure 2:**
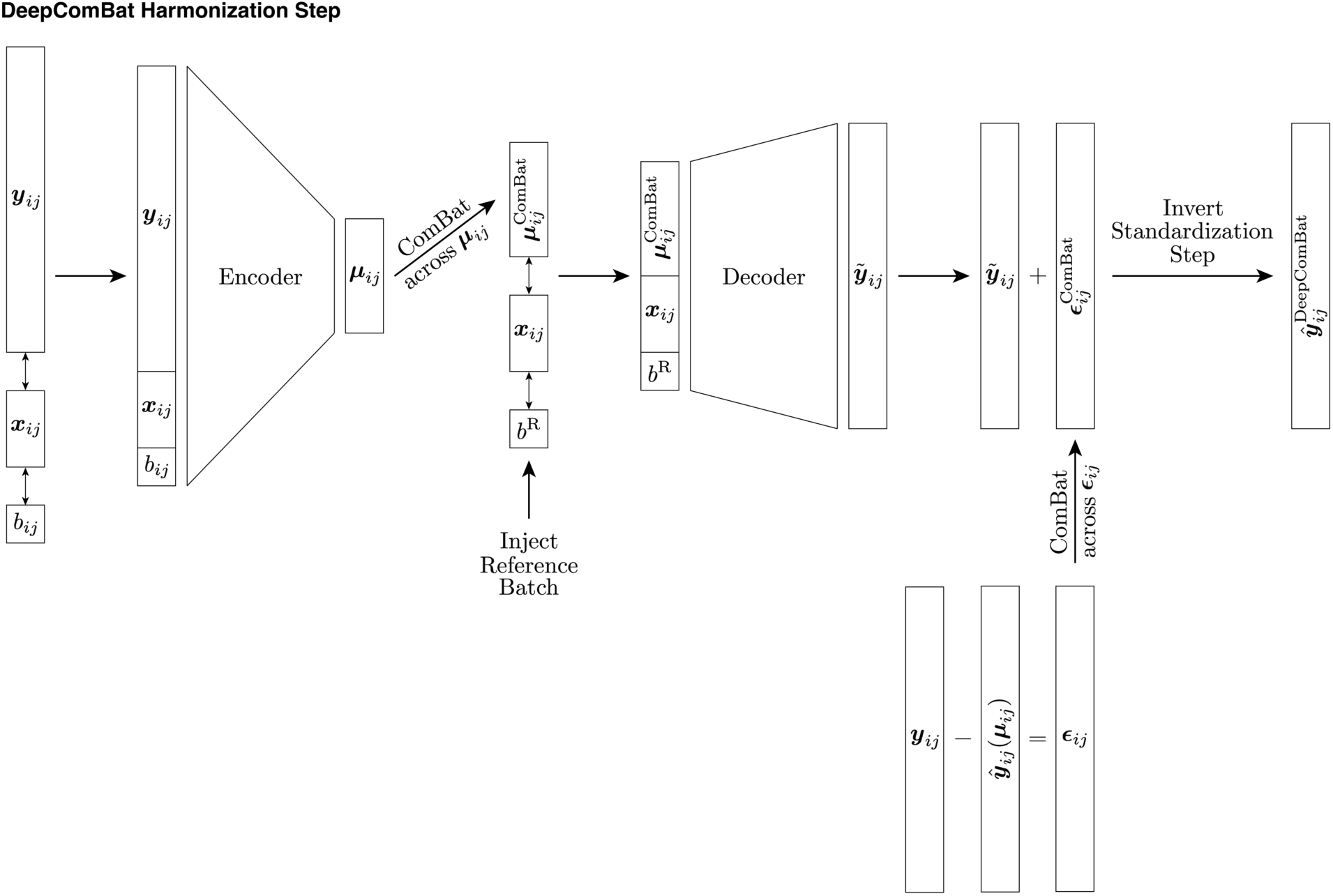
DeepComBat CVAE algorithm used during the harmonization step. At this step, encoder and decoder parameters have been learned during the training step and are frozen. Notation corresponds to that in the main text.

Next, the decoder output is harmonized. In this step, the decoder input is changed such that it receives harmonized latent space mean vectors as well as the desired batch for the harmonized data. The decoder additionally continues to receive unchanged biological covariates. Notationally, for one subject, this step can be represented as follows:

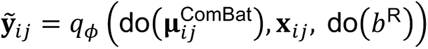

where 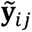 represents the harmonized decoder output, do(·), borrowed from the field of causal inference, represents the act of changing the decoder inputs, potentially contrary to fact, *b*^R^ is the desired reference batch for harmonization, and other notation is as defined above. Here, latent space distribution are changed to harmonized latent space mean vectors and batch is changed to the reference batch. Note that for all subjects, *b*^R^ must be the same such that all subjects are harmonized to the same batch, but that *b*^R^ can be defined to be either the first batch, the second batch, or some intermediate batch.

Then, the reconstruction residuals are calculated and harmonized. To estimate these residuals, noiseless reconstructions are first estimated by giving the decoder latent space mean vectors instead of the latent space distribution samples used during CVAE training. Notationally, for one subject, this is can be represented as:

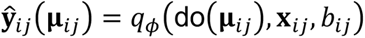

where 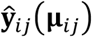 represents the noiseless reconstruction, in contrast to the noisy reconstruction used in training, 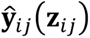.

Then, reconstruction residuals are defined as:

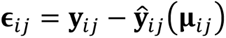

where ***y****_ij_* is the standardized raw data. These residuals are then corrected across all *ij* subjects using the ComBat model with both batch and biological covariates to obtain 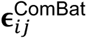.

Finally, individually harmonized components are combined and transformed back to the original feature space using the inverses of shift and scale parameters used to standardize the raw data in the normalization step:

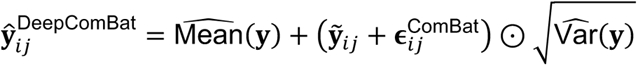

where 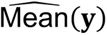 and 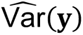 are the *p* × 1 vectors of overall feature-wise means and variances, respectively, used to standardize each feature in the normalization step, and ⊙ represents element-wise vector multiplication.

#### 2.3.6 Data and code availability

Data used in the preparation of this article were obtained from the Alzheimer’s Disease Neuroimaging Initiative (ADNI) database (adni.loni.usc.edu). The ADNI was launched in 2003 as a public-private partnership, led by Principal Investigator Michael W. Weiner, MD. The primary goal of ADNI has been to test whether serial magnetic resonance imaging (MRI), positron emission tomography (PET), other biological markers, and clinical and neuropsychological assessment can be combined to measure the progression of mild cognitive impairment (MCI) and early Alzheimer’s disease (AD). For up-to-date information, see www.adni-info.org.

An R package for performing DeepComBat is available at: https://github.com/hufengling/DeepComBat. This package is written in ‘torch for R’ which is the R analog of PyTorch that interfaces with the same C++ backend for fast computation. Across 30 runs, one full run of the DeepComBat algorithm on the ADNI dataset took an average of 53.0 seconds with standard deviation of 1.7 seconds on an Intel Xeon CPU with 2.40 GHz clock rate. Active memory use is negligible due to the mini-batch stochastic optimization routine.

The `deepcombat` package provides functions for normalization, DeepComBat architecture building, model-fitting, and manual hyperparameter tuning. It is designed to work on data in matrix form, where feature data to harmonize is stored in one matrix and biological and batch covariates are stored in another. Additionally, all code for evaluation and analysis is available at: https://github.com/hufengling/deepcombat_analyses.

Code for processing ADNI data is available at: https://github.com/ntustison/CrossLong.

### 2.4 Evaluation

DeepComBat was evaluated against unharmonized data as well as other feature-level harmonization methods where code was available. These methods included ComBat, CovBat, dcVAE, and gcVAE (An et al., 2022; Chen et al., 2022a; Fortin et al., 2017; Moyer et al., 2020). Notably, since no code was provided in the original manuscript for dcVAE, we implemented this method using code provided by An et al. (2022). For all comparison methods, we used default settings and hyperparameters provided in the code. Biological covariates of age, sex, and Alzheimer disease status were provided for ComBat, CovBat, and DeepComBat. Evaluation was conducted using qualitative visualization, statistical testing, and machine learning (ML) experiments.

In statistical testing and ML experiments, we assess the presence of both batch effects and biological effects. When assessing for batch effects, we assume that 1) test statistics corresponding to large p-values for statistical tests and 2) worse performance in predicting batch for ML experiments correspond to less presence of batch effects and therefore better harmonization. However, when assessing for biological effects, effective harmonization may lead to better, worse, or similar results, depending on the underlying relationship between batch and biological covariates.

On one hand, if batch status is strongly correlated with biological covariates in the dataset, removal of batch effects may decrease the magnitude of test statistics and/or predictive performance on biological covariates. On the other hand, if the presence of batch effects greatly reduces generalizability of models or the harmonization method introduces bias in the form of stronger biological covariate effects, the harmonization method may greatly increase the magnitude of test statistics and/or predictive performance on biological covariates. Finally, if none of these issues are present, harmonization may leave the resulting performance on biological covariates relatively unchanged, when compared to unharmonized data. In the ADNI dataset, based on the relative balance of biological covariates across batches, we assume that post-harmonization performance of both statistical testing and ML experiments should remain relatively unchanged. Therefore, we assess for biological effects mainly to investigate for potential pathological behavior rather than for direct comparison of harmonization methods.

#### 2.4.1 Qualitative visualization

We visualize the overall multivariate distribution of unharmonized and harmonized feature matrices using Unifold Manifold Approximation and Projection (UMAP) and principal component analysis (PCA) (McInnes et al., 2020). UMAP was fit using the `umap` package in R with 20 neighbors, 100 epochs, and default settings otherwise. Points were displayed by batch status. PCA was fit on correlation matrices to account for differences in scale across features. For UMAP and PCA, arbitrary differences in sign due to model fitting were changed in order to improve direct comparability of these visualizations between methods. Additionally, we explore how harmonization methods act on a small random sample of features using bivariate density plots and plots of feature-level changes after harmonization.

#### 2.4.2 Statistical testing

Harmonization methods were evaluated using mass univariate and multivariate statistical testing. For mass univariate testing, we performed a two-sample Anderson-Darling test on each feature, where the two samples were defined by batch covariate. Average p-value across all features and its standard deviation is reported. For this test, the null hypothesis is that the two samples come from the same distribution and the alternative hypothesis is that the two samples come from different distributions. Under the assumption that harmonization addresses distributional differences across batch, well-harmonized data should have non-significant p-values, and the mean p-value across all features should be approximately 0.5. Distributions of Anderson-Darling p-values across all features are also shown.

To test for differences in feature-wise means across batch as well as assess for validity of downstream analyses on biological covariates, we performed linear regression on each feature, where each regression model included the batch covariate as well as biological covariates of age, sex, and Alzheimer disease status. For each covariate, the average negative log 10 p-value across all features as well as the standard deviation of these transformed p-values is reported. Negative log 10 p-values are used to better represent the distribution of p-values very close to 0. Distributions of regression p-values for batch across all features are also shown.

For multivariate statistical testing, we assess harmonization results parametrically as well as non-parametrically. For parametric testing, we use the multivariate analysis of variance test (MANOVA), which tests for differences in multivariate means. The null hypothesis for this test is that there is no effect of a given covariate on the multivariate mean vector across all features while the alternative hypothesis is that there is some non-zero effect of the covariate on the multivariate mean vector. Our MANOVA model includes the batch covariate, age, sex, and Alzheimer disease status. We report the negative log 10 p-value based on Pillai’s trace test statistic, which has been shown to be more robust than other MANOVA test statistics (Olson, 1974).

For non-parametric multivariate testing, we use the k-nearest-neighbor batch-effect test (kBET) metric with default settings, developed and validated in the context of detecting batch effects in single-cell RNA-sequencing (scRNA-seq, Büttner et al., 2019). The kBET test is a non-parametric permutation-based test that 1) randomly samples a proportion of observations, 2) identifies each observation’s k-nearest neighbors, 3) evaluates whether the local distribution of batch among each set of k nearest neighbors differs from the global distribution of batch, and 4) generates an overall kBET statistic evaluating whether the number of observations with large differences in local distribution of batch are greater than that expected to occur by chance alone. Ultimately, the null hypothesis tested by kBET is that the observed local distributions of batch are similar to the expected local distributions of batch, conditional on the global distribution of batch.

#### 2.4.3 Machine learning experiments

To evaluate how our method interacts with multivariate batch or biological effects, we train ML algorithms to predict covariate information using the harmonized feature matrix. Prediction models were independently trained to perform classification of batch status, sex, and Alzheimer disease status, as well as regression of age. To perform the ML experiments, we use the `caret` package, version 6.0-93, to train and assess a large battery of ML algorithms on each feature matrix using the repeated cross-validation strategy, with five repeats of 10-fold cross-validation. This repeated cross-validation strategy was used to obtain a low-bias, low-variance estimate of the out-of-sample predictive performance.

For two-class classification of batch and sex, average area under the Receiver Operating Characteristic Curve (AUROC) across validation sets is reported. For three-class classification of Alzheimer disease status, AUROC cannot be calculated, so average accuracy across validation sets is reported. For regression of age, average *R*^2^ values across validation sets is reported. Note that in the repeated cross-validation strategy, average cross-validation metrics can be made arbitrarily precise by increasing the number of repeats, but variation in these metrics occurs within each cross-validation fold due to randomness in ML model fitting and train-validation splitting.

The ML evaluation battery for classification tasks consisted of: support vector machine (SVM) with radial basis, quadratic discriminant analysis (QDA), k-nearest neighbors (KNN), random forest (RF), and Extreme Gradient Boosted trees (XGBoost). The ML evaluation battery for regression of age consisted of: SVM with radial basis, KNN, RF, and XGBoost. SVM, QDA, KNN, and RF were fit using the default hyperparameters provided by their corresponding R packages. For XGBoost a few hyperparameters were *a priori* changed from the default to allow for greater algorithm differences when compared to RF. These changed hyperparameters included: `eta` = 0.1 and `colsample_bytree` = 0.5. Number of total boosting rounds had no default and was set to 100. Other hyperparameters were set to their defaults.

## 3 Results

### 3.1 DeepComBat reduces batch effects in qualitative visualizations

We visualized the effect of DeepComBat univariately and multivariately. For a representative, randomly-sampled region’s cortical thickness, density plots by batch revealed differences in distribution across batch in the raw data that could be attributed to differences in mean, variance, and shape (Figure 3). This distributional difference was qualitatively mitigated by ComBat, CovBat and DeepComBat, while dcVAE and gcVAE showed substantial transformation of the feature distribution.

**Figure 3:**
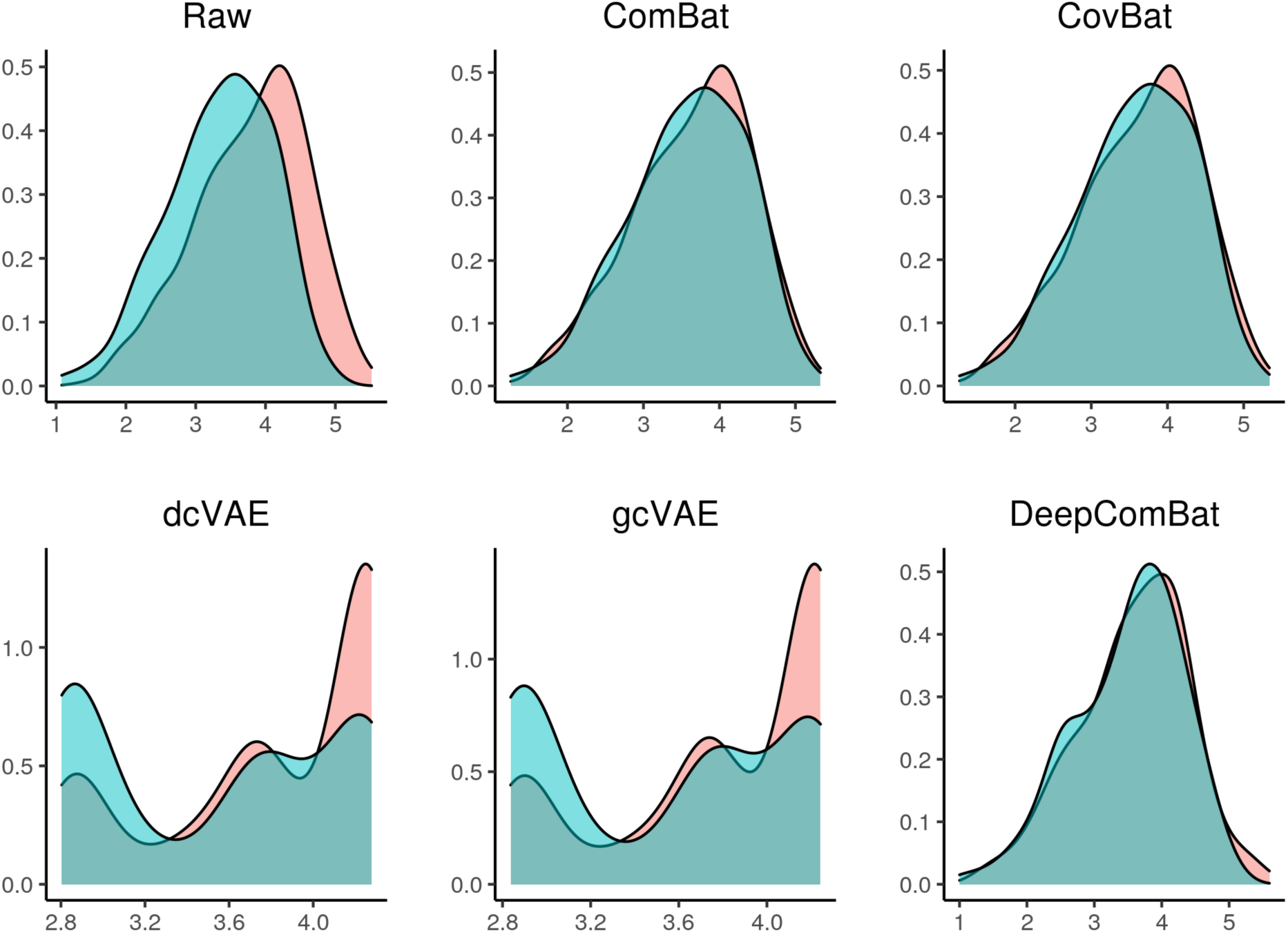
Density plots of one randomly sampled feature for raw data and various harmonization method outputs. Red color corresponds to the Siemens batch and blue color corresponds to the non-Siemens batch.

Similar qualitative results were observed in visualizations of the multivariate feature distribution using the first two UMAP dimensions and the first two principal components (Figures 4 and 5).

**Figure 4:**
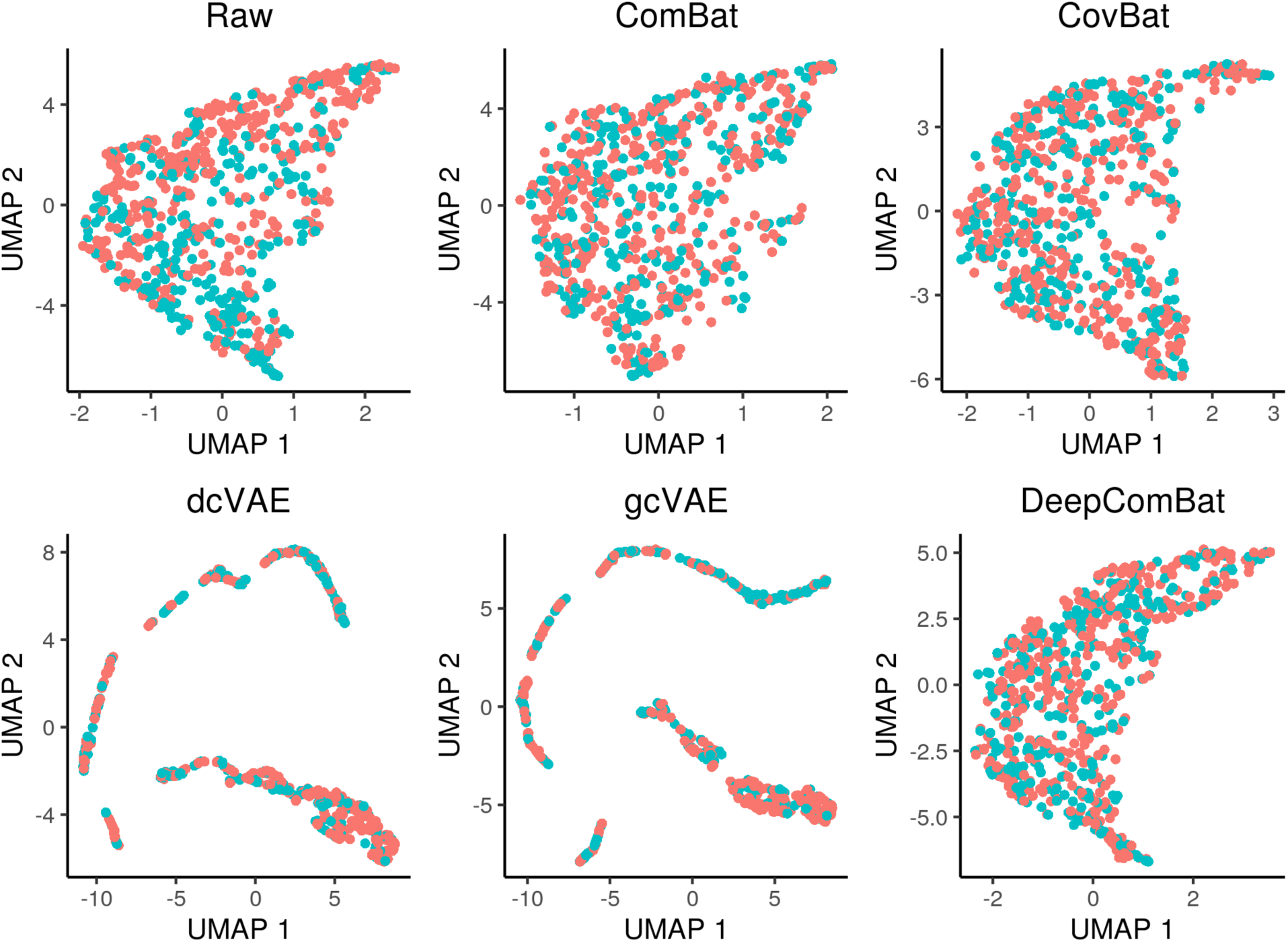
UMAP visualization of raw data and various harmonization method outputs. Red color corresponds to the Siemens batch and blue color corresponds to the non-Siemens batch.

**Figure 5:**
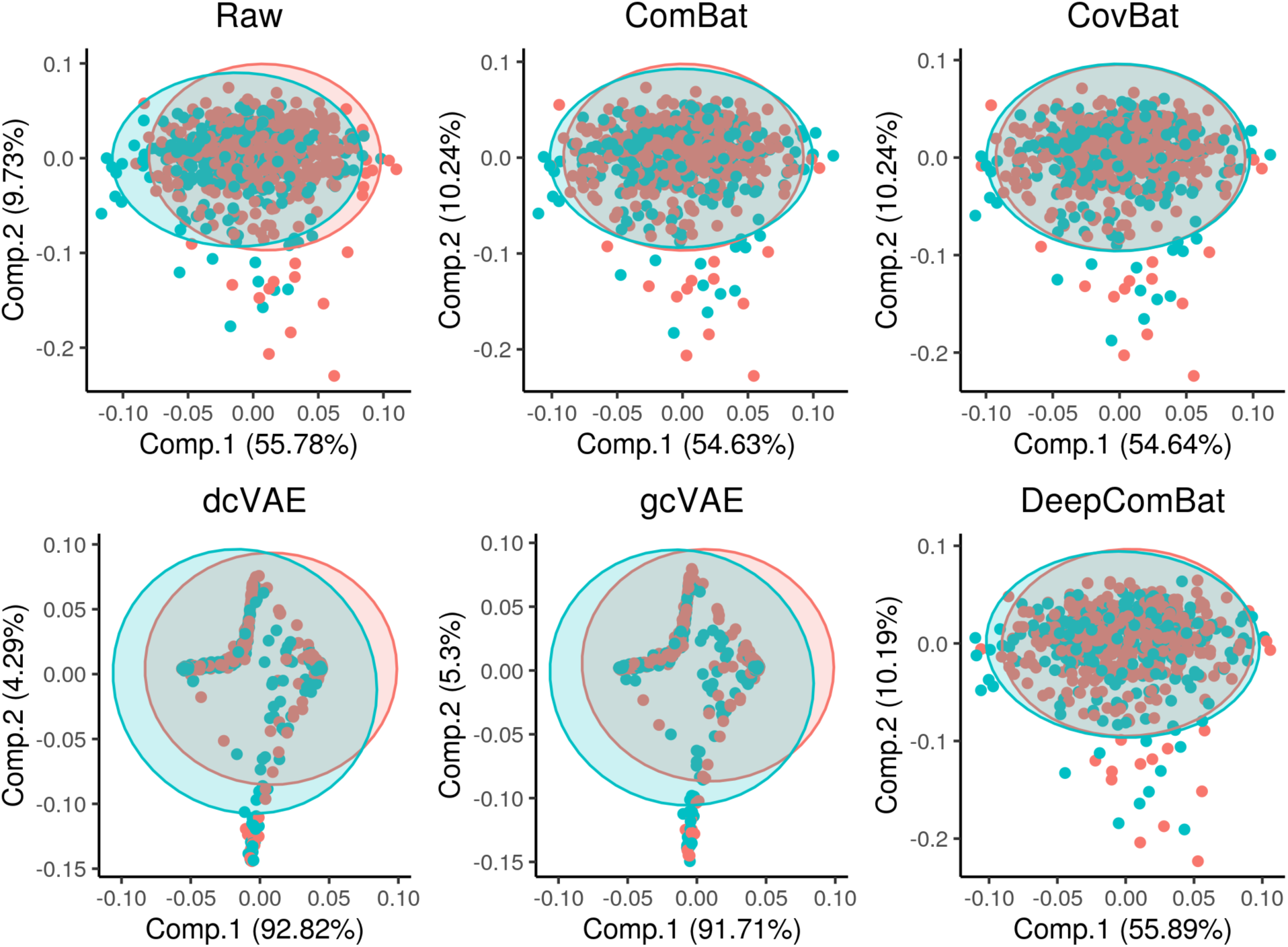
PCA visualization of raw data and various harmonization method outputs. PCA ellipses denote major and minor axes for each batch, centered at the batch-wise mean. Red color corresponds to the Siemens batch and blue color corresponds to the non-Siemens batch.

Finally, we explored how various harmonization methods change the raw data at the feature level. Here, we randomly sampled 10 cortical thickness features and randomly sampled 100 subjects to obtain a total of 1,000 randomly-sampled cortical thickness values. For each harmonization method, we plotted harmonized values for these cortical thicknesses against their corresponding raw values in Figure 6. In this visualization, ComBat and CovBat seemed to mostly induce upward and downward linear shifts in the data with small deviations from these shifts, which is consistent with their underlying shift and scale models. DeepComBat induced small non-linear shifts in the data on a similar scale as ComBat and CovBat. Meanwhile, dcVAE and gcVAE mapped harmonized values to their corresponding CVAE-predicted mean values without accounting for unmodeled CVAE reconstruction errors. Thus, dcVAE and gcVAE produced outputs with noise patterns characteristic of synthetic data, as noted by Dewey et al. (2019).

**Figure 6:**
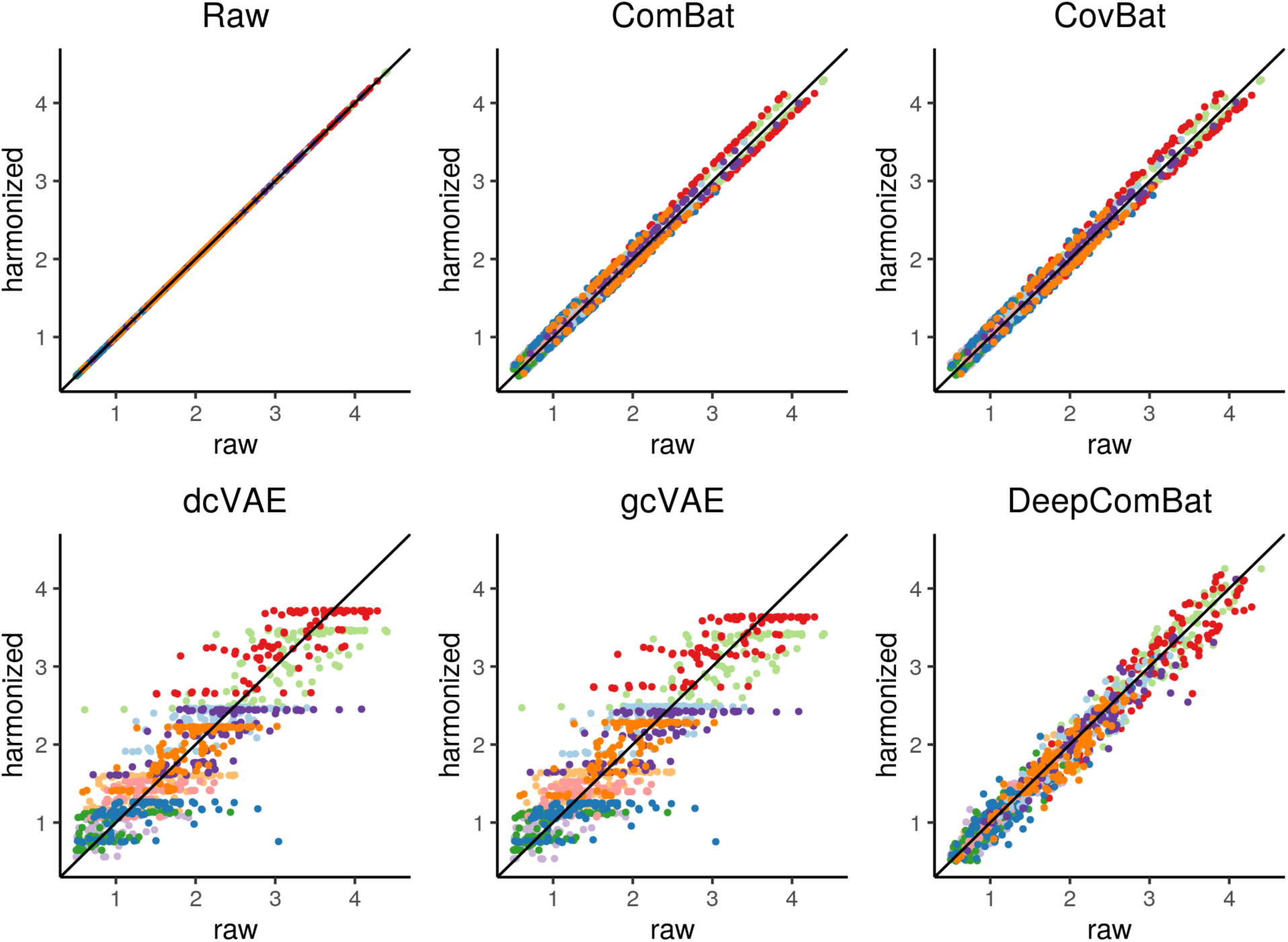
Visualization of randomly-sampled harmonized values plotted against their corresponding raw values for various harmonization methods. Colors indicate each of the 10 randomly-sampled cortical thickness features.

Overall, qualitative visualizations showed DeepComBat seemed to make reasonable changes to univariate feature distributions, preserve the underlying multivariate structure of the data, and estimate harmonized values that were highly correlated with the corresponding raw values.

### 3.2 DeepComBat removes statistically-detectable batch effects and preserves inference on biological effects

Average feature-wise Anderson-Darling test results, presented in Table 2, suggested there were significant differences in univariate distributions across batches in the raw data, which is consistent with the qualitative results seen in Figure 3. These differences were effectively reduced by ComBat, CovBat, and DeepComBat – each of these harmonization methods produced Anderson-Darling p-values with means around 0.5 and large standard deviations. However, dcVAE and gcVAE produced outputs such that Anderson-Darling p-values for all features were 0, suggesting large differences in distribution post-harmonization across features. These results are further illustrated in Figure 7, which presents quantile-quantile plots comparing observed negative log 10 feature-wise p-values to expected p-values under a uniform distribution. In this Figure, ComBat, CovBat, and DeepComBat show p-value distributions qualitatively similar to a uniform distribution, while raw data, dcVAE, and gcVAE showed p-value distributions with many more highly-significant p-values than expected under a uniform distribution.

**Figure 7:**
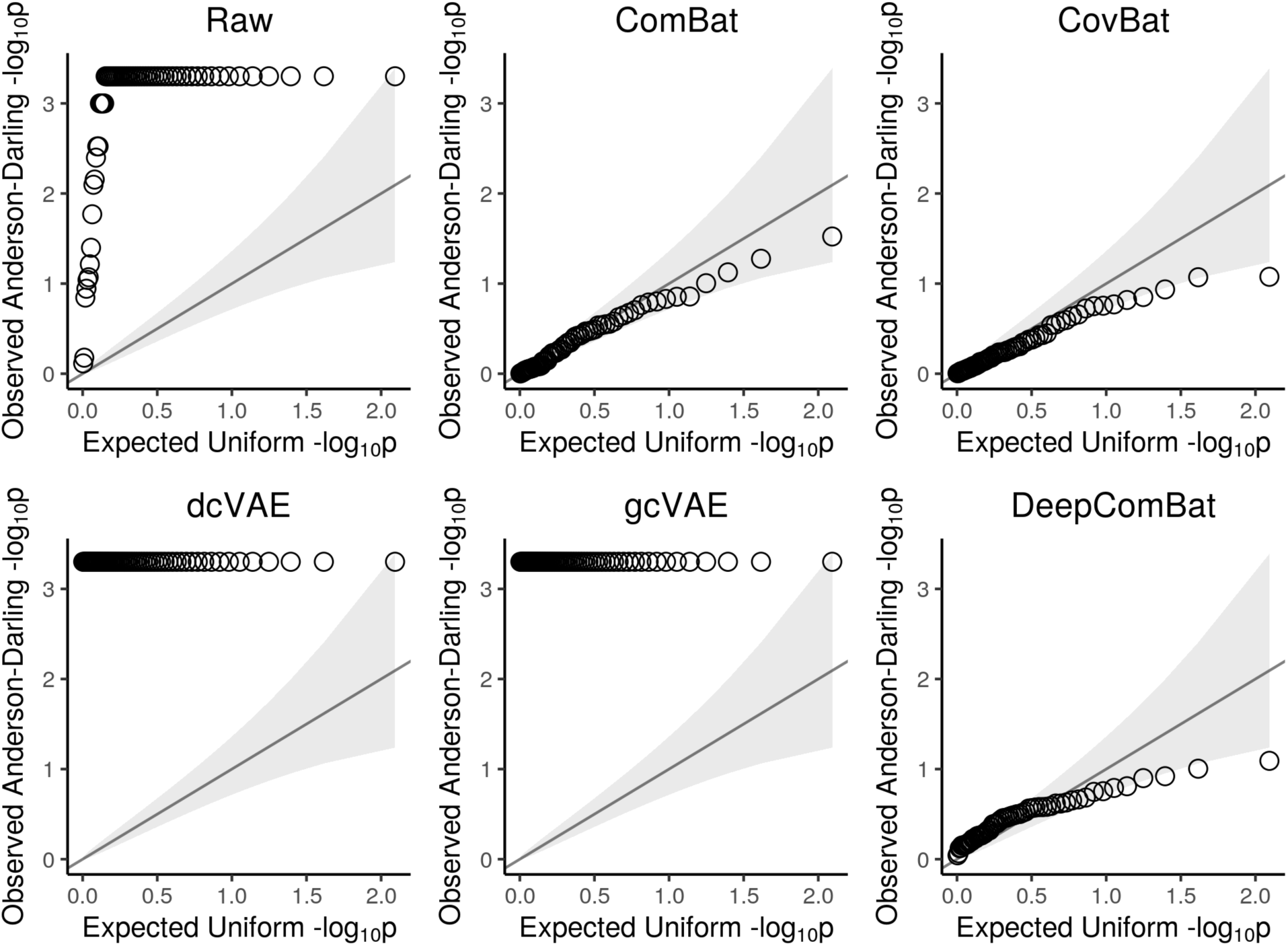
Quantile-Quantile (Q-Q) plots of observed feature-wise Anderson-Darling negative log 10 p-values from raw and harmonized data. Observed p-values are plotted against expected negative log 10 p-values under a uniform distribution. Gray band corresponds to 95% confidence intervals for whether observed data was sampled from a uniform.

**Table 2:** Average feature-wise Anderson-Darling p-values for batch.

|  | **Mean (SD)** |
| --- | --- |
| Raw | 0.03 (0.13) |
| ComBat | 0.52 (0.29) |
| CovBat | 0.56 (0.27) |
| dcVAE | 0 (0) |
| gcVAE | 0 (0) |
| DeepComBat | 0.42 (0.21) |

Feature-wise linear regression results are presented in Table 3 as average negative log 10 p-values and in Figure 8 as quantile-quantile plots of negative log 10 p-value distributions. In the table, p-values of 1, 0.05, and 0.01 correspond to negative log 10 p-values of 0, 1.3, and 2, respectively, with small p-values corresponding to large negative log 10 p-values. As with the Anderson-Darling analysis, this analysis on the raw data also showed significant differences in mean across batch, when age, sex, and Alzheimer disease status were also included in the model. Differences in batch-wise means were effectively removed by ComBat, CovBat, and DeepComBat, while dcVAE and gcVAE seemed to increase the difference in batch-wise means. As in the Anderson-Darling results, DeepComBat seemed to provide slightly less correction of univariate batch differences when compared to CovBat, but provided similar levels of correction when compared to ComBat. Similar results are qualitatively observed in Figure 8.

**Figure 8:**
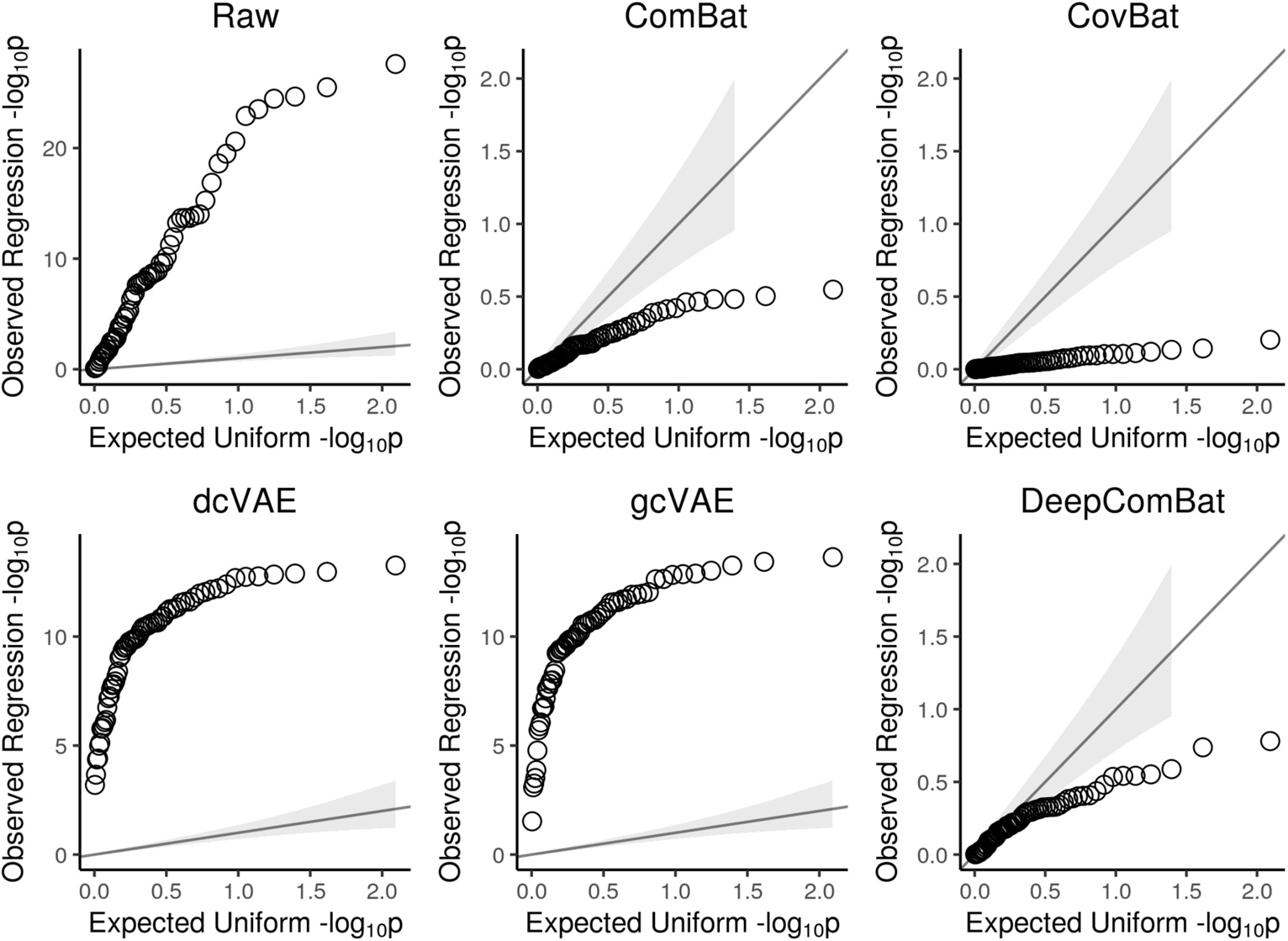
Quantile-Quantile (Q-Q) plots of observed feature-wise linear regression negative log 10 p-values from raw and harmonized data. Observed p-values are plotted against expected negative log 10 p-values under a uniform distribution. Gray band corresponds to 95% confidence intervals for whether observed data was sampled from a uniform. Y-axis scales differ between panels.

**Table 3:** Average feature-wise linear regression results. Reported as negative log 10 p-values - Mean (SD). Negative log 10 of conventional p-value threshold 0.05 is 1.30. Larger is more significant.

|  | Age | Sex | AD Status (CN) | AD Status (LMCI) | Batch |
| --- | --- | --- | --- | --- | --- |
| Raw | 15.08 (6.25) | 0.83 (0.76) | 15.51 (7.75) | 6.02 (3.11) | 8.7 (7.44) |
| ComBat | 15.14 (6.27) | 0.83 (0.76) | 15.55 (7.77) | 6.02 (3.12) | 0.19 (0.15) |
| CovBat | 15.14 (6.27) | 0.84 (0.76) | 15.56 (7.8) | 6.05 (3.14) | 0.05 (0.04) |
| dcVAE | 23.68 (3.07) | 0.41 (0.1) | 26.7 (2.89) | 9.27 (1.81) | 9.48 (2.63) |
| gcVAE | 23.46 (3.87) | 0.41 (0.11) | 26 (3.69) | 9.01 (1.83) | 9.48 (2.86) |
| DeepComBat | 14.99 (6.31) | 0.85 (0.78) | 16.16 (8.24) | 6.18 (3.11) | 0.25 (0.18) |

Additionally, ComBat, CovBat, and DeepComBat preserved inference on biological covariates of age, sex, and Alzheimer disease status, with the distribution of corresponding negative log 10 p-values sharing similar means and standard deviations when compared to that of the raw data. Notably, DeepComBat was seen to slightly increase power for detecting average differences between controls and both LMCI and AD subjects in cortical thicknesses. Meanwhile, dcVAE and gcVAE showed large increases in power for age and Alzheimer disease effects and a large decrease in power for sex effects. These large increases in statistical power for age and Alzheimer disease effects may be explained by the exclusion of unmodeled residuals in dcVAE and gcVAE harmonized outputs.

In statistical testing for multivariate effects using MANOVA, ComBat, CovBat, and DeepComBat were seen to completely remove batch effects from the multivariate mean across features when biological covariates were also included (Table 4). dcVAE and gcVAE were also able to provide a substantial degree of multivariate batch effects correction when compared to the raw data; however, the MANOVA p-value still remained highly significant indicating significant batch effects remained.

**Table 4:** Multivariate analysis of variance (MANOVA) results. Reported as negative log 10 p-values. Negaive log 10 of conventional p-value threshold 0.05 is 1.30. Larger is more significant.

|  | **Age** | **Sex** | **AD Status** | **Batch** |
| --- | --- | --- | --- | --- |
| Raw | 33.56 | 25.50 | 15.55 | 101.73 |
| ComBat | 31.26 | 22.14 | 15.20 | 0.00 |
| CovBat | 31.27 | 22.01 | 15.35 | 0.00 |
| dcVAE | 9.30 | 0.46 | 10.40 | 11.96 |
| gcVAE | 13.20 | 0.43 | 11.72 | 10.63 |
| DeepComBat | 32.66 | 18.54 | 17.65 | 0.00 |

As in the feature-wise linear regression analysis, ComBat, CovBat, and DeepComBat were additionally able to preserve inference on biological covariates. Notably, DeepComBat was slightly less powerful for multivariate sex effects and slightly more powerful for Alzheimer disease effects when compared to the raw data. The decrease in power for sex effects may reflect removal of the confounding between sex and batch seen in Table 1 while the increase in power for inference on Alzheimer disease effects may reflect removal of batch-attributable noise.

Finally, in non-parametric testing using kBET, CovBat and DeepComBat were able to produce harmonized outputs where the distributions of batch within local neighborhoods were not significantly different from the global distribution of batch. In contrast, kBET detected highly significant differences in local distributions of batch for raw data, ComBat, dcVAE, and gcVAE; the proportion of local neighborhoods with detectable differences in batch distributions was much less in ComBat compared to all of raw data, dcVAE and gcVAE.

**Table 5:**
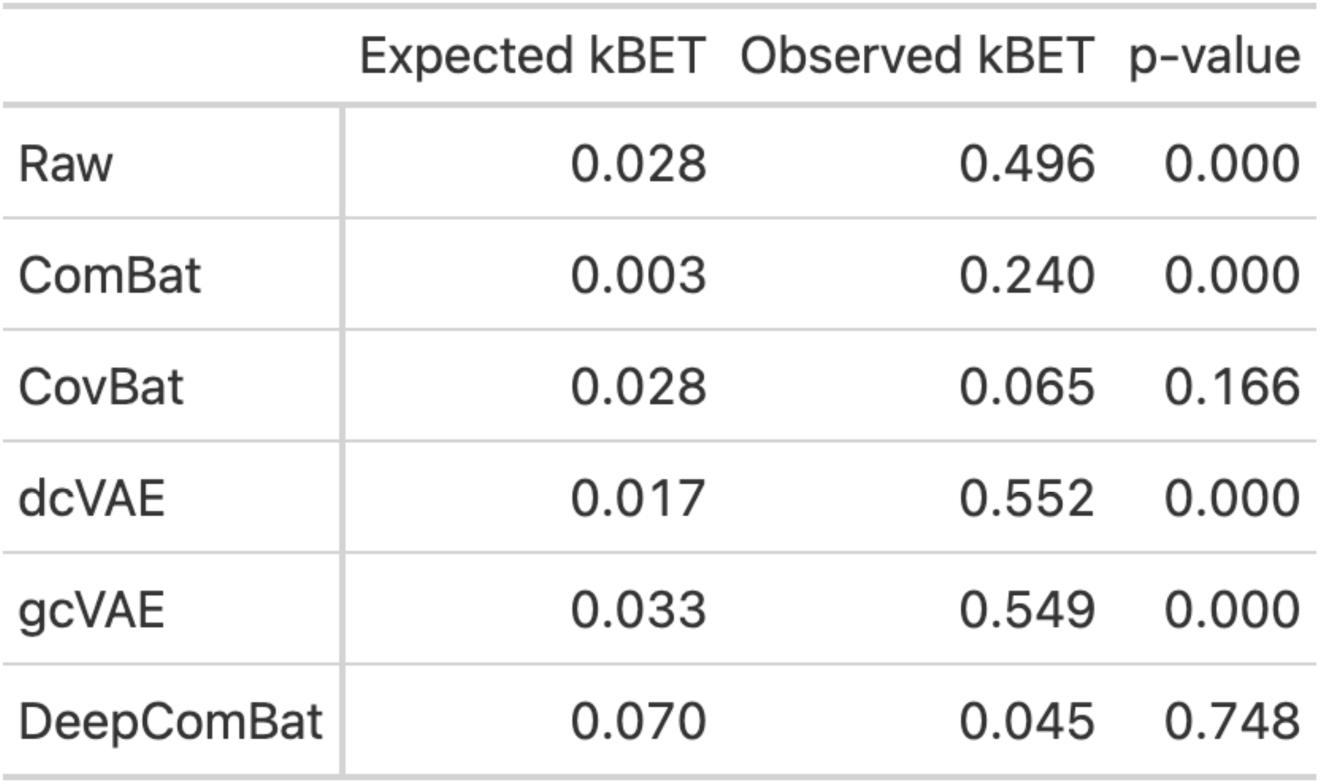
kBET results for batch.

|  | Expected kBET | Observed kBET | p-value |
| --- | --- | --- | --- |
| Raw | 0.028 | 0.496 | 0.000 |
| ComBat | 0.003 | 0.240 | 0.000 |
| CovBat | 0.028 | 0.065 | 0.166 |
| dcVAE | 0.017 | 0.552 | 0.000 |
| gcVAE | 0.033 | 0.549 | 0.000 |
| DeepComBat | 0.070 | 0.045 | 0.748 |

Overall, we found that DeepComBat effectively removes statistically-detectable batch effects both univariately and multivariately, and in doing so, is also able to effectively preserve biological information without introducing bias from the statistical-inference perspective. In univariate analyses, DeepComBat performed slightly worse than ComBat and CovBat in terms of harmonization, while in multivariate analyses, DeepComBat outperformed ComBat and CovBat in terms of harmonization. Regarding preserving and increasing power for detecting biological associations, DeepComBat performed favorably compared to ComBat and CovBat. DeepComBat outperformed dcVAE and gcVAE by all metrics.

### 3.3 DeepComBat impairs detection of batch by ML algorithms and maintains predictability of biological covariates

A battery of ML experiments seeking to predict batch status were run on raw and harmonized data (Figure 9). Note that the error bars shown in this section represent the standard deviations, not standard errors of the mean.

**Figure 9:**
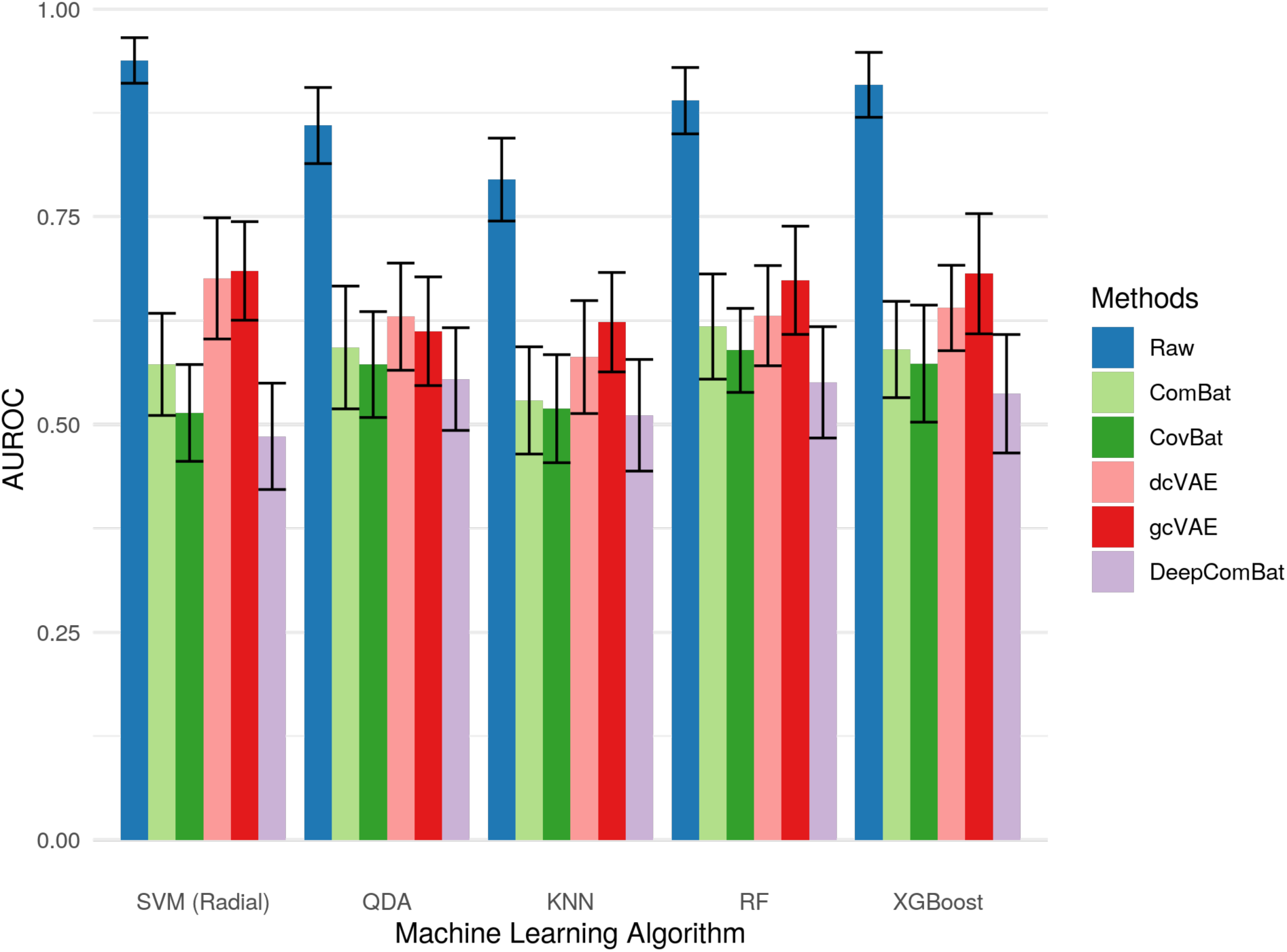
Bar graphs showing average AUROC for predicting batch of various classifiers on raw and harmonized data. Error bars represent the standard deviation of validation-set AUROCs across 5 repeats of 10-fold cross-validation.

All classifiers could effectively determine the batch status of out-of-sample subjects in the raw data. This ability to detect batch was greatly decreased by all harmonization methods, with DeepComBat-harmonized data consistently corresponding to the lowest AUROCs across all ML experiments. CovBat-harmonized data corresponded to the second lowest AUROCs.

Additionally, DeepComBat effectively retained biological information in its outputs. In Figures 10, 11, and 12, DeepComBat-harmonized data showed predictive performances similar to, or better than that, of raw, ComBat-corrected, and CovBat-corrected data. All post-harmonization predictive performances were significantly higher than those of dcVAE and gcVAE.

**Figure 10:**
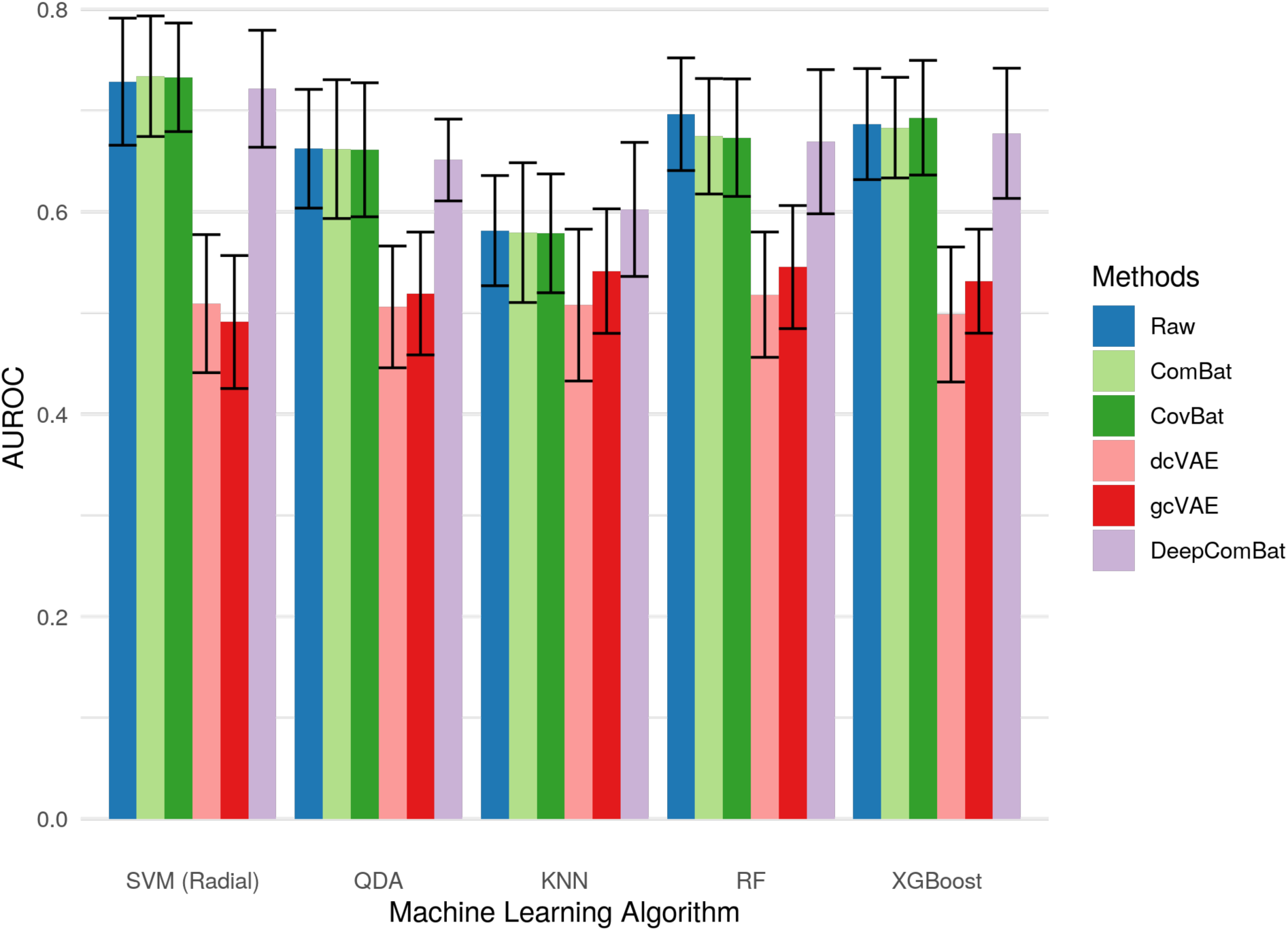
Bar graphs showing average AUROC for predicting sex of various classifiers on raw and harmonized data. Error bars represent the standard deviation of validation-set AUROCs across 5 repeats of 10-fold cross-validation.

**Figure 11:**
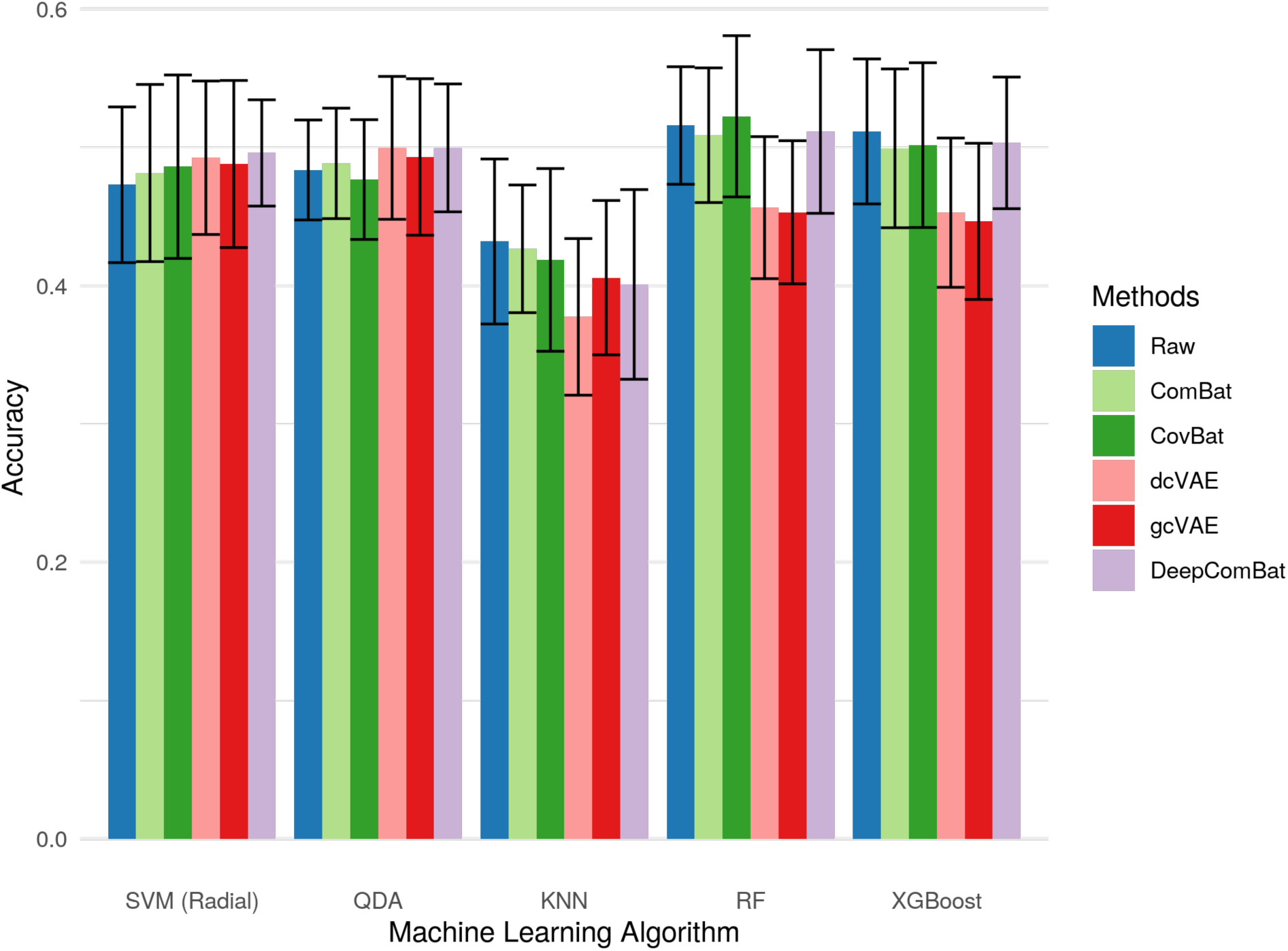
Bar graphs showing average accuracy for predicting Alzheimer disease status of various classifiers on raw and harmonized data. Error bars represent the standard deviation of validation-set accuracies across 5 repeats of 10-fold cross-validation.

**Figure 12:**
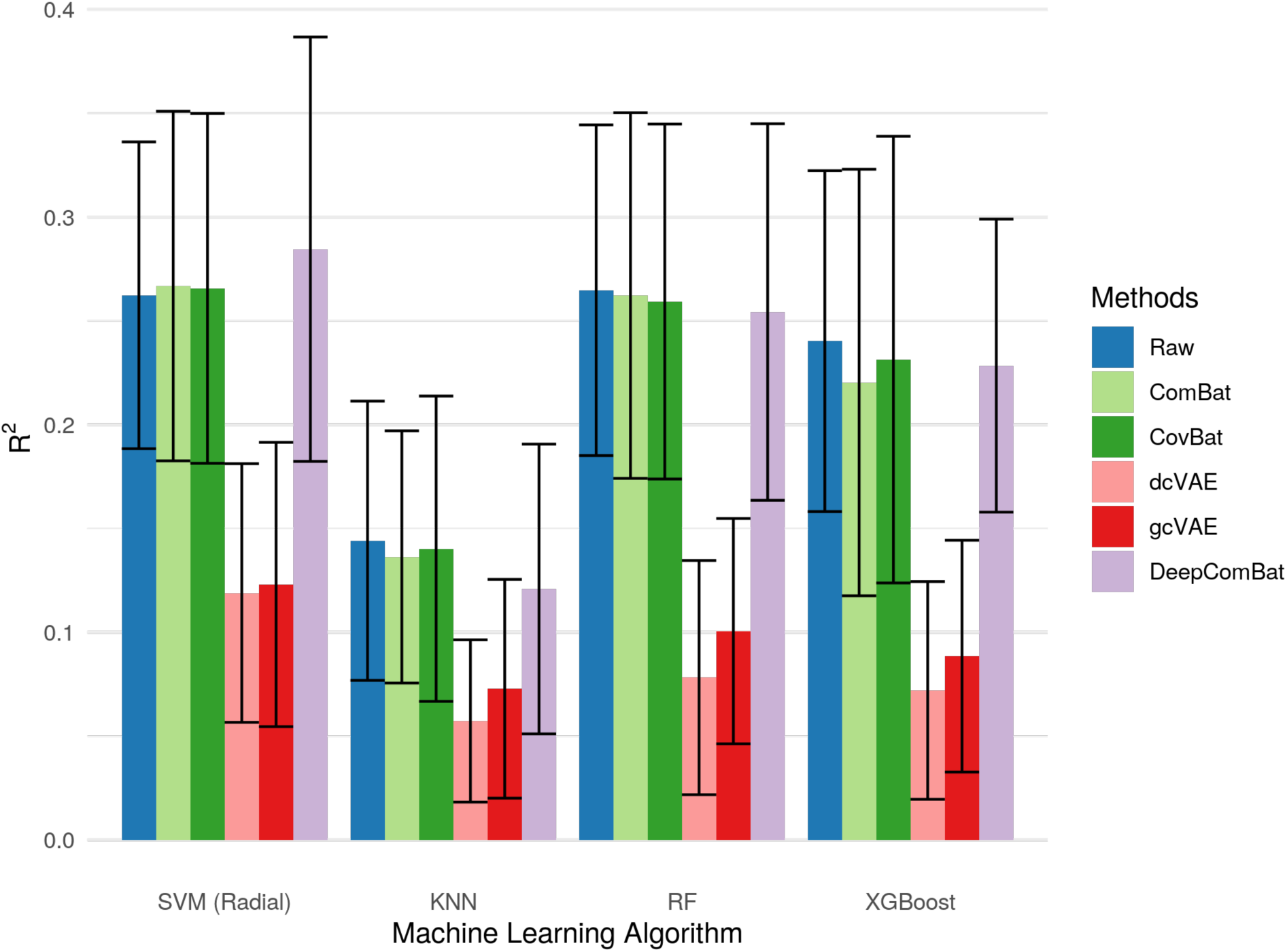
Bar graphs showing average *R*^2^ value for predicting age of various classifiers on raw and harmonized data. Error bars represent the standard deviation of validation-set *R*^2^ values across 5 repeats of 10-fold cross-validation.

Overall, we found that DeepComBat more effectively removed multivariate batch effects than other harmonization methods, even when assessed via powerful ML algorithms such as XGBoost. Additionally, DeepComBat effectively preserved biological information in the predictive context.

### 3.4 DeepComBat is easily tunable and robust to selection of the KL-divergence weighting hyperparameter

We find the DeepComBat KL-divergence hyperparameter, *λ*_Final_, can be easily tuned manually through investigation of the variances of latent space distributions since *λ*_Final_ is directly correlated with these variances. Empirically, we find that a good choice of *λ*_Final_ can be found when the overall distribution of natural-log variances across all latent space dimensions is qualitatively similar to the one shown in the *λ* = 0.1 panel of Figure 13. Namely, it is desirable that the distribution of log variances is bimodal with some latent-space dimensions being informative (those with very negative log variances) and some latent-space dimensions being non-informative (those with log variances near 0).

**Figure 13:**
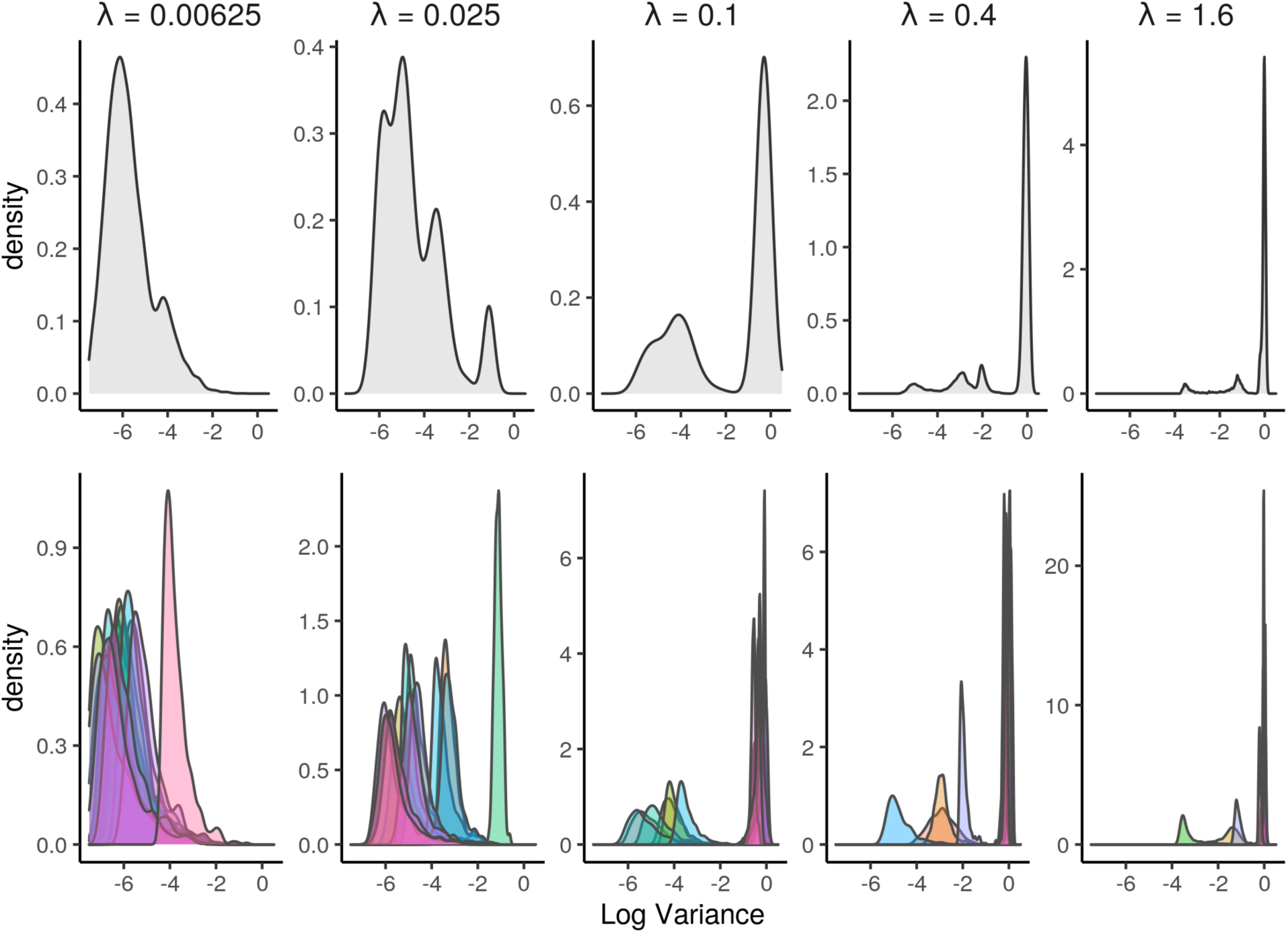
Top: Density plots of the log variances of latent-space distributions for each subject over all latent-space dimensions for a range of hyperparameter choices. Bottom: Density plots of the log variances of latent-space distributions for each subject for individual latent-space dimensions for a range of hyperparameter choices.

If too many latent-space dimensions are informative, as in the first two panels of Figure 13, *λ*_Final_ should be increased, as DeepComBat may leave too many batch effects in the latent space. If too many latent-space dimensions are uninformative, as in the last two panels of Figure 13, *λ*_Final_ should be decreased, as DeepComBat will be unable to adequately reconstruct the subject-specific mean, and batch effects will be left in the CVAE residuals. Thus, examination of log variance density plots allows for easy heuristics for choosing a suitable hyperparameter value.

However, even when *λ*_Final_ is empirically misspecificed, DeepComBat performs effective harmonization. The above analyses, including qualitative visualizations, statistical tests, and ML experiments, were run on DeepComBat-harmonized outputs produced using the range of choices for *λ*_Final_ ranging from 16 times greater than the *λ*_Final_ used in our primary analysis to values 16 times less. These robustness analyses are presented in Supplementary Tables 1-4 and Supplementary Figures 1-9.

Notably, results demonstrated comparable DeepComBat performance across all evaluations for these choices of *λ*_Final_. Moreover, results from the DeepComBat-harmonized output corresponding to *λ*_Final_ = 0.4 were superior to the results from our primary analysis. This suggests that more intensive hyperparameter tuning may further improve harmonization performance, though this improvements comes at the cost of additional complexity in model fitting, computation time, and user expertise.

This robustness result may be due to the design of DeepComBat, which partitions batch effects originally present in the raw data into one of these three components: the CVAE latent space, the CVAE decoder, and the reconstructed residuals. Component-wise harmonization therefore allows for a form of “double-robustness” with respect to CVAE fitting. In worst-case scenarios, if *λ*_Final_ is too large such that most batch effects are contained in the reconstruction residuals, ComBat on these residuals will still allow for reasonable overall harmonization; and if *λ*_Final_ is too small such that most batch effects are contained in the latent space, ComBat on the latent space will address the batch effects. Thus, DeepComBat is robust to misspecification of *λ*_Final_ – as long as the specified *λ*_Final_ is reasonably close to the optimal KL-divergence weight, final DeepComBat output data will be sufficiently harmonized.

## 4 Discussion

Multi-batch neuroimaging data are increasingly common and necessary for learning generalizable models for inference and prediction. There is also growing interest in using ML techniques to perform multivariate pattern analysis and train powerful classifiers that can efficiently use multivariate data. To enable these efforts, there is increasing need for statistically-rigorous multivariate harmonization methods.

In this study, we demonstrate that strong batch effects exist in raw data, and that these batch effects remain detectable by ML experiments even after state-of-the-art statistical harmonization methods are applied. We also find that, while previously-proposed deep learning harmonization approaches are able to partially remove batch effects from the ADNI dataset, this batch effects correction comes at the cost of removal of relevant biological information as well as introduction of artifacts characteristic of synthetic data. We then propose DeepComBat, a novel hybrid method that is able to take advantage of the strengths of both deep learning and statistical methods – it uses the CVAE architecture to perform non-linear, multivariate correction as well as the ComBat model to rigorously and robustly harmonize the latent space and residuals. When compared to other methods, we show DeepComBat performs more effectively when evaluated by highly-multivariate ML experiments as well as non-parametric kBET testing. It performs comparably to ComBat and CovBat when evaluated by statistical tests. Overall, these results suggest that DeepComBat may be especially useful for harmonization in settings where prediction or inference using multivariate features and multivariate methods is the goal. In these settings, feature-wise correction using statistical methods may lead to significant non-corrected batch effects that may be picked up by prediction methods and inappropriately used.

### 4.1 DeepComBat may be more robust to model misspecification when compared to statistical methods

Similarly to many statistical methods, such as ComBat and CovBat, DeepComBat assumes batch effects can be estimated as differences in feature-wise conditional means and variances of unmodeled residuals. However, unlike statistical methods, mean batch effect are estimated non-linearly and multivariately using a combination of batch and biological covariates along with subject-specific latent space representations. In this mean batch effect estimation procedure, mean batch effect are partially removed by the decoder in a non-parametric manner, where the only assumption on the nature of batch effect is that it can be approximated by the decoder network. Thus, while latent space harmonization still involves the ComBat model, overall harmonization may be less contingent on how well the data follow ComBat assumptions. Additionally, discussed further below, moment-matching of latent space representations has been empirically shown to be effective in various harmonization-like tasks (Fatania et al., 2022; Huang and Belongie, 2017; Lopez et al., 2018; Lotfollahi et al., 2019; Zuo et al., 2021).

In terms of correcting batch effect in unmodeled residuals, DeepComBat argues a meaningful portion of what statistical methods claims are “unmodeled residuals” – information that is not explained by biological nor batch covariates by the naive linear model – can in fact be explained as a multivariate non-linear function of biological covariates, batch covariates, and subject-specific latent factors. Through the CVAE architecture, DeepComBat is able to significantly reduce the mean squared error between model-predicted feature vectors and raw feature vectors when compared to ComBat and CovBat. Thus, DeepComBat is able to directly model and correct more batch effect in terms of conditional differences in mean, and less batch effect is corrected based on the strong assumption that there are batch-wise differences in the variances of unmodeled residuals. Subsequently, although DeepComBat still uses the ComBat model to correct the residuals, it may rely less on ComBat-specific assumptions since the magnitude of batch effect correction on the residuals is smaller.

### 4.2 DeepComBat relaxes strong assumptions made by other deep learning methods and simplifies model fitting

Previous feature-level deep learning harmonization methods, including dcVAE and gcVAE make a number of strong implicit assumptions. These assumptions include 1) perfect model fit, which assumes that reconstruction residuals insignificant and therefore do not need to accounted for or re-incorporated, 2) fully disentangleable latent space, which assumes that the neural network can completely learn a batch-invariant latent space based on the loss function alone, and 3) balanced biological covariates across batches, which assumes that all population-level differences across batch are in fact due to batch and should be removed.

However, the first assumption is violated in situations where the latent-space dimensions are too small to adequately capture non-batch information, the decoder is not complex enough to efficiently encode all batch-related information, and sample sizes within batches are too limited to estimate all the necessary network parameters. These violations are further compounded when the first two assumptions are considered together; while near-perfect model fit may be achievable with a large latent space, it is even more challenging when a completely batch-invariant latent space is required. Finally, in neuroimaging datasets where biological covariates are imbalanced across batches, such as in the ADNI dataset used in this study, complete removal of marginal batch-wise differences will necessarily involve removal of biological information as well.

DeepComBat is able to relax these strong implicit assumptions by 1) accounting for the presence of reconstruction residuals and re-introducing them on top of the CVAE-harmonized subject-level means, 2) explicitly removing batch effects from the CVAE latent space, and 3) conditioning on biological covariates at each harmonization step. By relaxing these assumptions, we are able to greatly improve the usability of DeepComBat by simplifying its architecture when compared to that of dcVAE and gcVAE. For example, dcVAE and gcVAE rely on adversarial training with a discriminator in order to train their decoders to produce more realistic outputs, but DeepComBat no longer needs this adversarial component since non-perfect model fit is acceptable. This minimizes computational burden and avoids common challenges in adversarial training. DeepComBat also circumvents the need for precise tuning of the KL divergence weighting hyperparameter, since remaining batch effects in the latent space are explicitly removed after CVAE training.

Importantly, relaxing these assumptions allows for DeepComBat to be designed such that, if a subject-level feature vector is “self-harmonized” back to its actual batch, that feature vector will be unchanged. This makes sense, since “self-harmonization” should be the identity function. However, in other deep learning harmonization methods, including dcVAE and gcVAE, since reconstruction residuals are not explicitly accounted for in these other methods, the “self-harmonized” data will have less noise. This phenomena has been highlighted by Dewey et al. (2019), who noticed that DeepHarmony, an image-level harmonization method, produced harmonized images with noise characteristics indicative of a synthetic image – namely, that they looked smoother. By keeping reconstruction residuals in the final harmonized output, DeepComBat avoids an implicit assumption of perfect model fit and allows for harmonized outputs to retain natural noise characteristics.

### 4.3 DeepComBat resembles other moment-matching harmonization methods

DeepComBat primarily achieves multivariate harmonization by using ComBat in the CVAE latent space in order to generate a batch-invariant latent space. In this step, ComBat is used as a moment-matching model that takes advantage of shrinkage estimation in order to match conditional means and variances across batches. Analogies between latent-space ComBat and other moment-matching style transfer algorithms can be drawn.

Specifically, in single-cell RNA-sequencing (scRNA-seq) batch effects correction, scGen encodes gene expression data to a latent space using a standard variational autoencoder (Lotfollahi et al., 2019). Then, the algorithm estimates and removes mean batch effects, or first moments, from this latent space, conditional on cell type, in order to perform correction. CVAE-based methods such as dcVAE, gcVAE, and a number of scRNA-seq methods, such as scVI, can also be thought to perform latent-space moment-matching (Lopez et al., 2018); however, these methods do so implicitly through the loss function, rather than by explicitly estimating and correcting latent space coordinates.

Additionally, in image style transfer, where the goal is to change the style of an image without changing its content, adaptive instance normalization (AdaIN) can be used along with a convolutional autoencoder and its variations (Huang and Belongie, 2017). In the convolutional autoencoder, images are encoded into a set of latent space convolutional filters. Then, AdaIN performs style transfer by matching the means and variances of each filter, learned from the original image, to the means and variances of the corresponding filters learned from the image that has the desired style. In the non-convolutional setting of DeepComBat, each 1 × 1 element of the latent space vector corresponds to one convolutional filter, and similar moment-matching is performed, but at the group level instead of the individual input level.

Finally, outside of deep learning methods, DeepComBat draws on ideas from CovBat, which has been shown to harmonize the mean and covariance across sites. CovBat first performs standard ComBat and then corrects the covariance structure of residuals by projecting them into a latent space defined by principal components and running ComBat again. Thus, CovBat performs univariate mean harmonization and linearly-multivariate residual harmonization. DeepComBat flips these steps – it first performs non-linear multivariate mean harmonization and then univariately corrects the reconstruction residuals. Notably, DeepComBat autoencoder residuals are much smaller in magnitude than CovBat linear model residuals, so univariate residual correction is sufficient.

### 4.4 Limitations and future directions

DeepComBat is designed to only require minimal hyperparameter tuning, and this design choices improve the usability of DeepComBat by end-users that are not deep learning experts. Even so, DeepComBat performance can vary across different choices of hyperparameter as well as across different training runs of the same hyperparameter choice. Notably, DeepComBat still performs effective harmonization across a range of hyperparameter choices and training runs; however, stochastic outputs from a complex algorithm may be undesirable for end-users seeking deterministic and transparent harmonization behavior.

While DeepComBat is intended to train quickly on standard computing resources, such as laptops, DeepComBat is still much slower than statistical methods. The overall training time is further increased when manual hyperparameter tuning is taken into consideration, as end-users may need to train a few models before choosing a suitable *λ*_Final_. Overall, hyperparameter tuning and final model training should take no longer than 5-10 minutes, depending on the number of hyperparameters tried and dataset size.

Relatedly, we show DeepComBat performs better than dcVAE and gcVAE, which were trained using the default hyperparameter values specified by An et al. (2022). These default hyperparameter values were determined based on a different dataset, so they may not have been optimal for our ADNI data. Hyperparameter tuning for these two methods by thorough grid search may have improved their results; however, the computational and coding difficulty associated with hyperparameter tuning reflect challenges in applying these methods and sensitivity to hyperparameter misspecification. Further work could involve incorporating automated hyperparameter selection for these methods as well as DeepComBat in order to provide more optimal outputs. Notably, while better hyperparameter choices may have improved dcVAE and gcVAE performance, these methods still implicitly make the assumptions described above.

Next, as a deep learning model, DeepComBat may be reliant on relatively large sample sizes to appropriately estimate neural network parameters. In this study, we showed DeepComBat can effectively harmonize a dataset of 663 individuals across two batches; however, in datasets with smaller sample sizes where batch effects are harder to precisely estimate, purely statistical methods such as ComBat and CovBat may perform better. In this same vein, a strength of DeepComBat is in its ability to project high-dimensional feature data into a low-dimensional latent space that is easier to harmonize – datasets with fewer baseline features may benefit less from this approach. Finally, DeepComBat may be unnecessarily complex when intended downstream analyses involve using features in a univariate manner. For example, standard ComBat may be sufficient if feature-wise inference is the goal, but if the objective is to train a ML classifier, DeepComBat may be more appropriate.

Finally, although DeepComBat partially uses neural networks to learn and correct non-linear, multivariate batch effects without the need for explicitly specifying an underlying model, overall harmonization still requires the ComBat model in the latent space and residual space. Thus, standard ComBat limitations still apply. For example, information about biological covariates not conditioned on in DeepComBat may be removed along with batch effects, and non-linear covariate effects located within the latent space or residuals may be inappropriately estimated and removed along with batch effects. Further work could consider relaxing these limitations by allowing for complex, non-linear covariate effects in the latent space and residuals (Pomponio et al., 2020). Additionally, like standard ComBat, DeepComBat assumes independence between subjects and is thus designed for cross-sectional data – extensions to longitudinal data may be an important next step (Beer et al., 2020).

## 5 Conclusion

DeepComBat is a novel, statistically-rigorous, deep learning approach to image harmonization that leverages deep learning and statistical concepts to perform multivariate batch effects correction conditional on biological covariates. We demonstrate it can more effectively remove multivariate batch effects from structural neuroimaging feature while preserving biological information than previously-proposed methods. As high-dimensional, multi-batch data becomes more common and interest in using ML techniques to analyze such data grows, we hope that DeepComBat will serve as a tool for end-users to remove multivariate batch confounding as well as provide a new perspective for methodologists to develop improved harmonization methods.

## Acknowledgements

This study was supported by grants from the National Institute of Neurological Disorders and Stroke (R01NS085211 and R01NS060910), the National Multiple Sclerosis Society (RG-1707-28586), and the University of Pennsylvania Center for Biomedical Image Computing and Analytics (CBICA). Funding sources were not involved in study design, data analysis, manuscript preparation, or submission decisions.

Data collection and sharing for this project was funded by the Alzheimer’s Disease Neuroimaging Initiative (ADNI) (National Institutes of Health Grant U01 AG024904) and DOD ADNI (Department of Defense award number W81XWH-12-2-0012). ADNI is funded by the National Institute on Aging, the National Institute of Biomedical Imaging and Bioengineering, and through generous contributions from the following: AbbVie, Alzheimer’s Association; Alzheimer’s Drug Discovery Foundation; Araclon Biotech; BioClinica, Inc.; Biogen; Bristol-Myers Squibb Company; CereSpir, Inc.; Cogstate; Eisai Inc.; Elan Pharmaceuticals, Inc.; Eli Lilly and Company; EuroImmun; F. Hoffmann-La Roche Ltd and its affiliated company Genentech, Inc.; Fujirebio; GE Healthcare; IXICO Ltd.; Janssen Alzheimer Immunotherapy Research & Development, LLC.; Johnson & Johnson Pharmaceutical Research & Development LLC.; Lumosity; Lundbeck; Merck & Co., Inc.; Meso Scale Diagnostics, LLC.; NeuroRx Research; Neurotrack Technologies; Novartis Pharmaceuticals Corporation; Pfizer Inc.; Piramal Imaging; Servier; Takeda Pharmaceutical Company; and Transition Therapeutics. The Canadian Institutes of Health Research is providing funds to support ADNI clinical sites in Canada. Private sector contributions are facilitated by the Foundation for the National Institutes of Health (www.fnih.org). The grantee organization is the Northern California Institute for Research and Education, and the study is coordinated by the Alzheimer’s Therapeutic Research Institute at the University of Southern California. ADNI data are disseminated by the Laboratory for Neuro Imaging at the University of Southern California.

