## Supplemental Tables and Figures for "DeepComBat: A Statistically Motivated, Hyperparameter-Robust, Deep Learning Approach to Harmonization of Neuroimaging Data"

### 7 Supplementary Materials

Supplementary Table 1: Average feature-wise Anderson-Darling p-values for batch.

| | Mean (SD) |
|---|---|
| Raw | 0.03 (0.13) |
| Lambda = 0.00625 | 0.41 (0.23) |
| Lambda = 0.025 | 0.45 (0.26) |
| Lambda = 0.1 | 0.42 (0.21) |
| Lambda = 0.4 | 0.46 (0.22) |
| Lambda = 1.6 | 0.44 (0.26) |

Supplementary Table 2: Average feature-wise linear regression results.

Reported as negative log 10 p-values — Mean (SD). Negative log 10 of conventional p-value threshold 0.05 is 1.30. Larger is more significant.

| | Age | Sex | AD Status (CN) | AD Status (LMCI) | Batch |
|---|---|---|---|---|---|
| Raw | 0.21 (0.18) | 0.55 (0.47) | 0.47 (0.48) | 0.34 (0.33) | 0.47 (0.36) |
| Lambda = 0.00625 | 15.02 (6.3) | 0.86 (0.79) | 16.18 (8.2) | 6.19 (3.15) | 0.26 (0.21) |
| Lambda = 0.025 | 15 (6.32) | 0.84 (0.77) | 16.01 (8.11) | 6.12 (3.06) | 0.27 (0.18) |
| Lambda = 0.1 | 14.99 (6.31) | 0.85 (0.78) | 16.16 (8.24) | 6.18 (3.11) | 0.25 (0.18) |
| Lambda = 0.4 | 15.08 (6.39) | 0.87 (0.81) | 16.36 (8.3) | 6.21 (3.13) | 0.2 (0.14) |
| Lambda = 1.6 | 15.23 (6.51) | 0.87 (0.79) | 16.46 (8.32) | 6.29 (3.17) | 0.28 (0.18) |

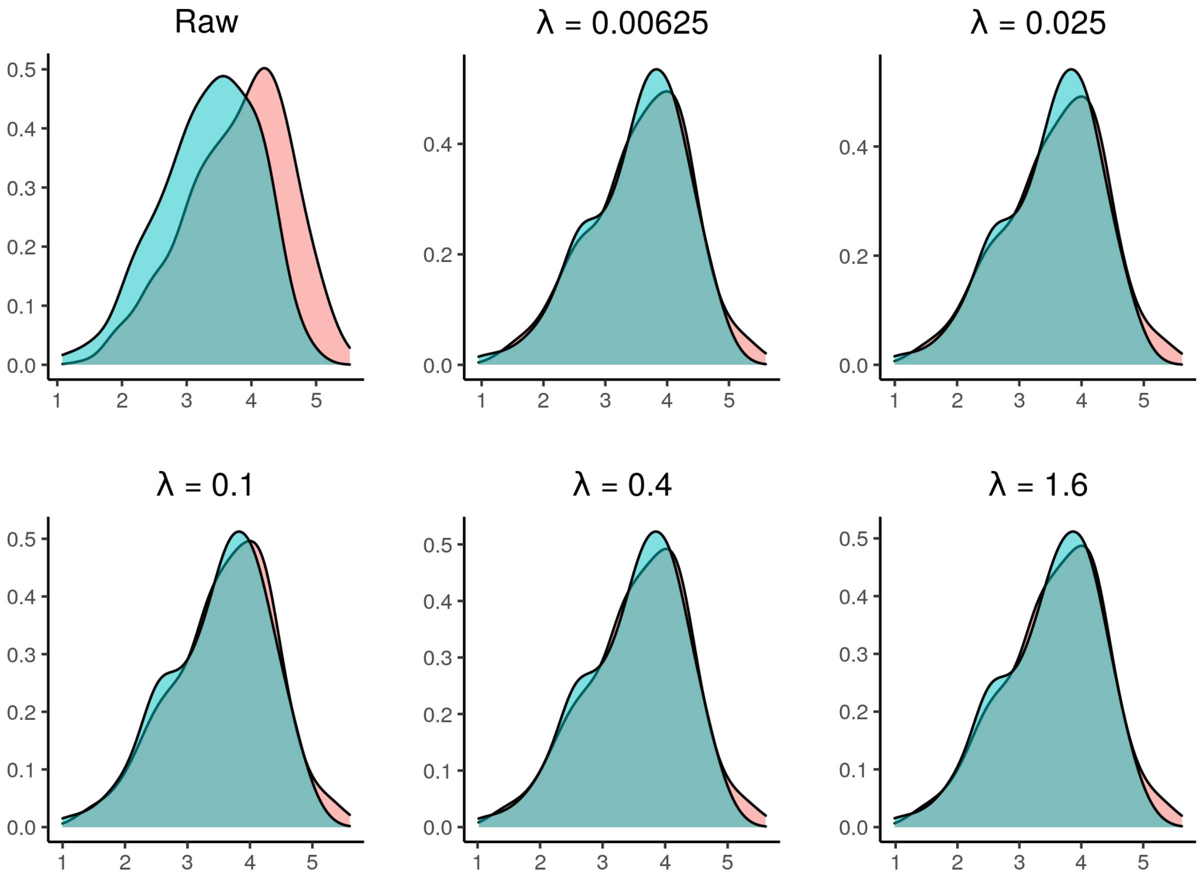


*Supplementary Figure 1: Density plots of one randomly sampled feature for raw data and harmonized data across different hyperparameter choices. Red color corresponds to the Siemens batch and blue color corresponds to the non-Siemens batch.*

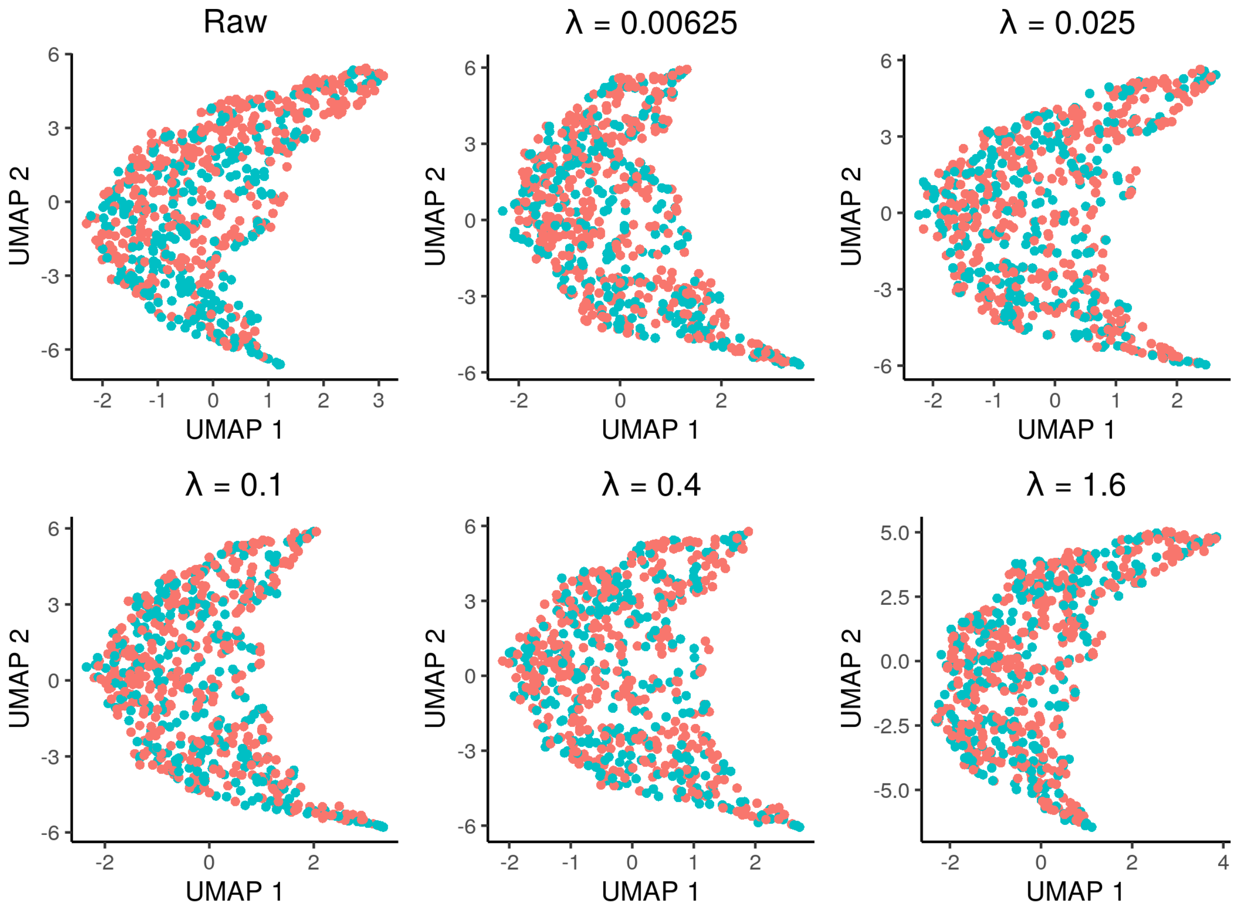


*Supplementary Figure 2: UMAP visualization of raw data and harmonized data across different hyperparameter choices. Red color corresponds to the Siemens batch and blue color corresponds to the non-Siemens batch.*

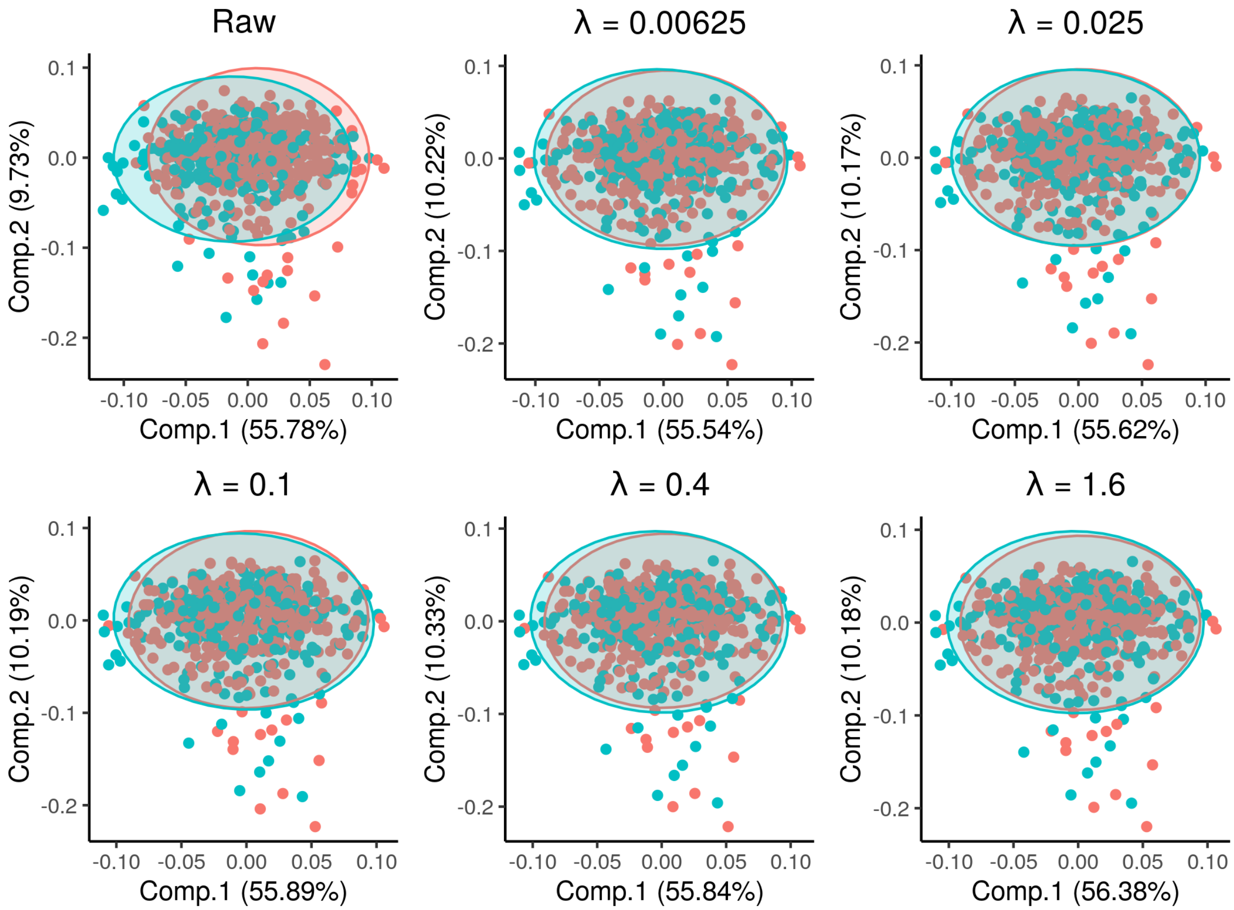


*Supplementary Figure 3: PCA visualization of raw data and harmonized data across different hyperparameter choices. PCA ellipses denote major and minor axes for each batch, centered at the batch-wise mean. Red color corresponds to the Siemens batch and blue color corresponds to the non-Siemens batch.*

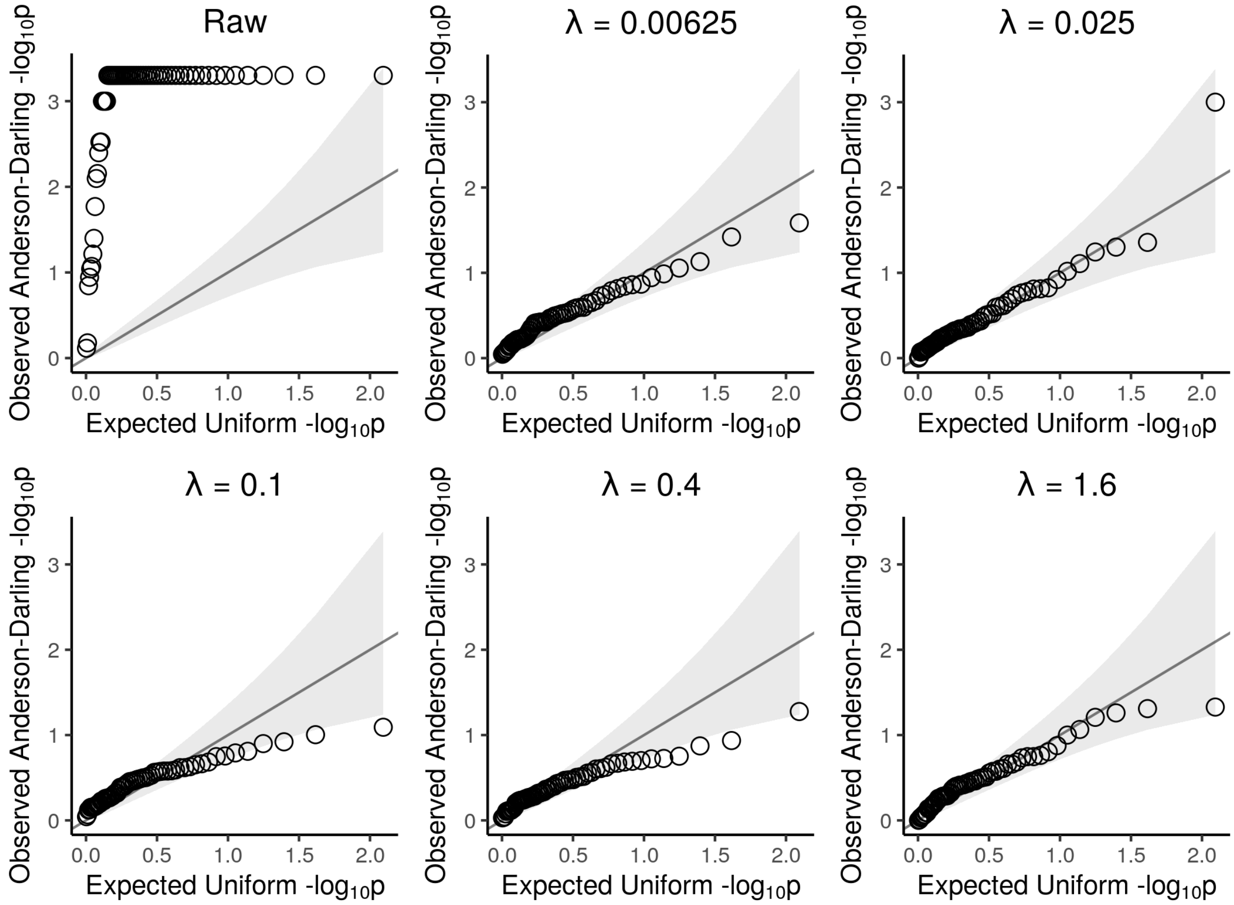


*Supplementary Figure 4: Quantile-Quantile (Q-Q) plots of observed feature-wise Anderson-Darling negative log 10 p-values from raw data and harmonized data across different hyperparameter choices. Observed p-values are plotted against expected negative log 10 p-values under a uniform distribution. Gray band corresponds to 95% confidence intervals for whether observed data was sampled from a uniform.*

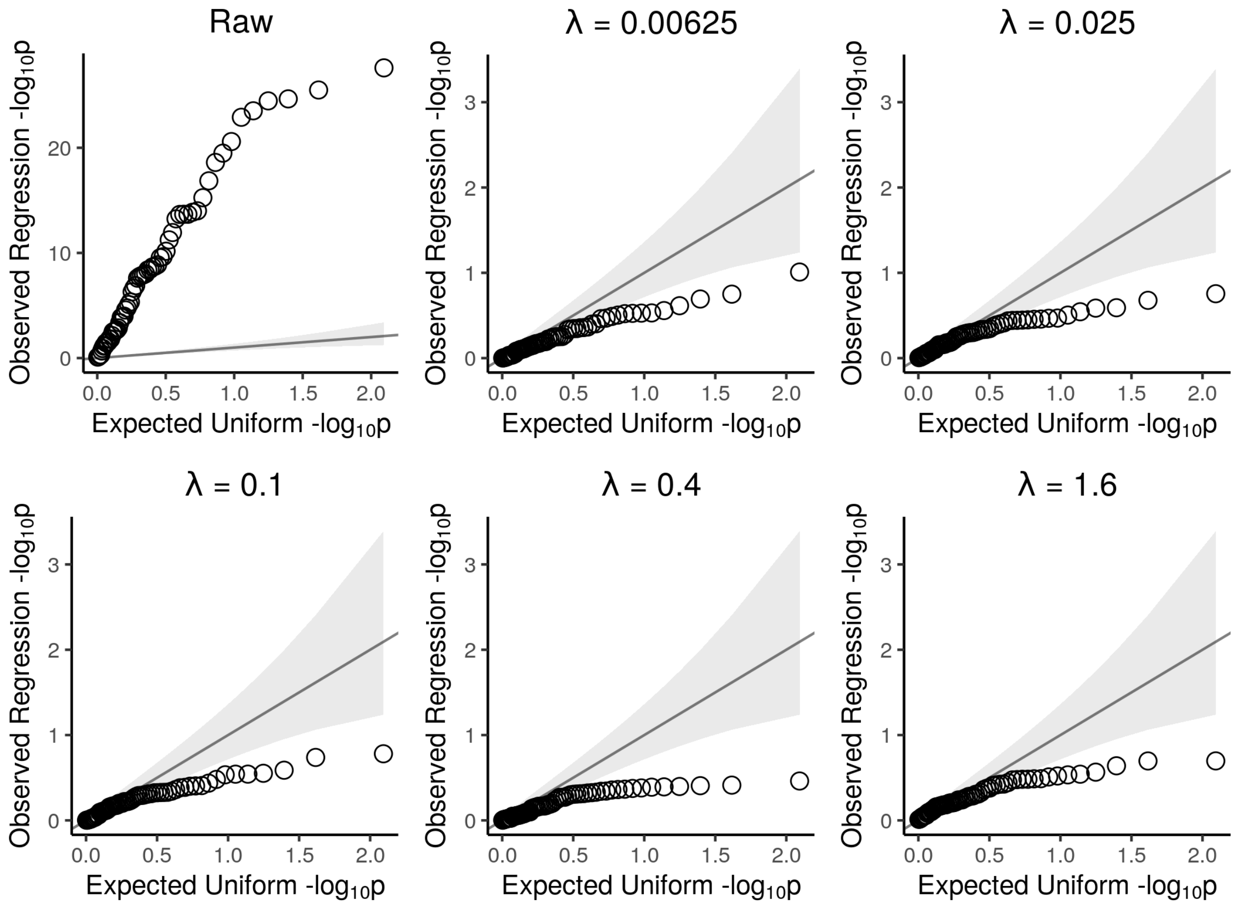


*Supplementary Figure 5: Quantile-Quantile (Q-Q) plots of observed feature-wise linear regression negative log 10 p-values from raw data and harmonized data across different hyperparameter choices. Observed p-values are plotted against expected negative log 10 p-values under a uniform distribution. Gray band corresponds to 95% confidence intervals for whether observed data was sampled from a uniform. Y-axis scales differ between panels.*

#### Supplementary Table 3: Multivariate analysis of variance (MANOVA) results.

Reported as negative log 10 p-values. Negative log 10 of conventional p-value threshold 0.05 is 1.30. Larger is more significant.

| | Age | Sex | AD Status | Batch |
|---|---|---|---|---|
| Raw | 33.56 | 25.50 | 15.55 | 101.73 |
| Lambda = 0.00625 | 34.02 | 18.91 | 17.90 | 0.00 |
| Lambda = 0.025 | 33.38 | 17.77 | 17.40 | 0.00 |
| Lambda = 0.1 | 32.66 | 18.54 | 17.65 | 0.00 |
| Lambda = 0.4 | 33.94 | 19.05 | 18.30 | 0.00 |
| Lambda = 1.6 | 34.35 | 18.72 | 18.17 | 0.00 |

Supplementary Table 4: kBET results for batch

| | Expected kBET | Observed kBET | p-value |
|---|---|---|---|
| Raw | 0.028 | 0.496 | 0.000 |
| Lambda = 0.00625 | 0.037 | 0.030 | 0.623 |
| Lambda = 0.025 | 0.022 | 0.038 | 0.344 |
| Lambda = 0.1 | 0.070 | 0.045 | 0.748 |
| Lambda = 0.4 | 0.015 | 0.035 | 0.289 |
| Lambda = 1.6 | 0.066 | 0.056 | 0.611 |

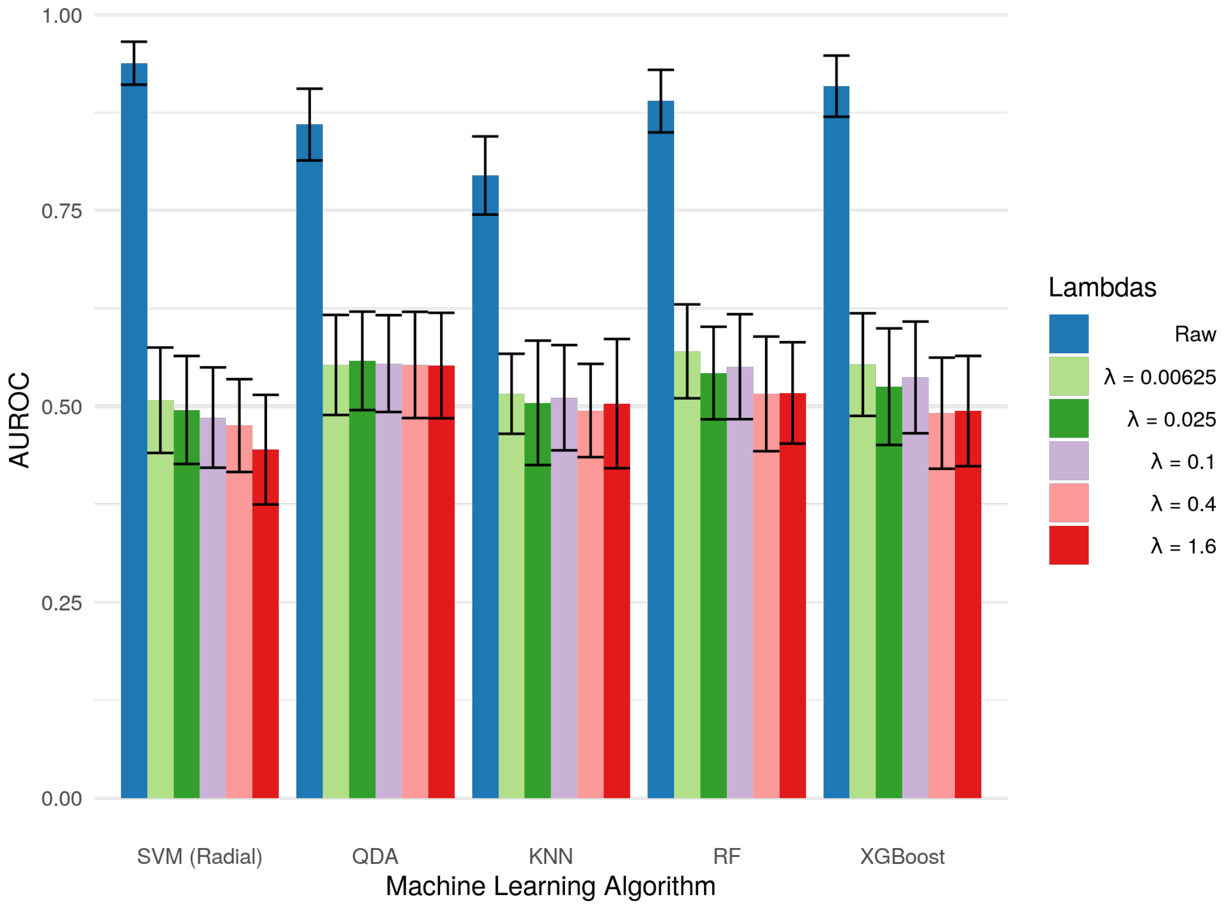


*Supplementary Figure 6: Bar graphs showing average AUROC for predicting batch of various classifiers on raw and harmonized data across different hyperparameter choices. Error bars represent the standard deviation of validation-set AUROCs across 5 repeats of 10-fold cross-validation.*

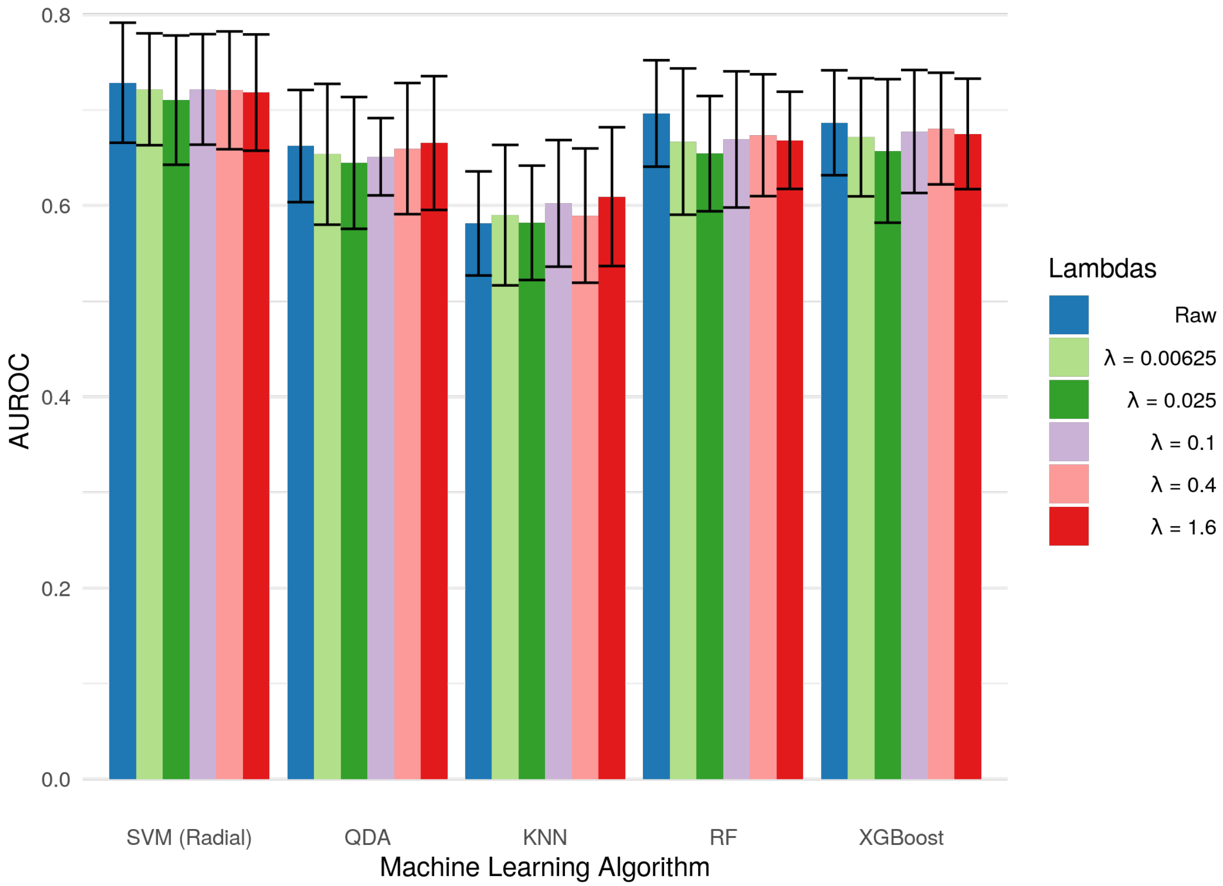


*Supplementary Figure 7: Bar graphs showing average AUROC for predicting sex of various classifiers on raw and harmonized data across different hyperparameter choices. Error bars represent the standard deviation of validation-set AUROCs across 5 repeats of 10-fold cross-validation.*

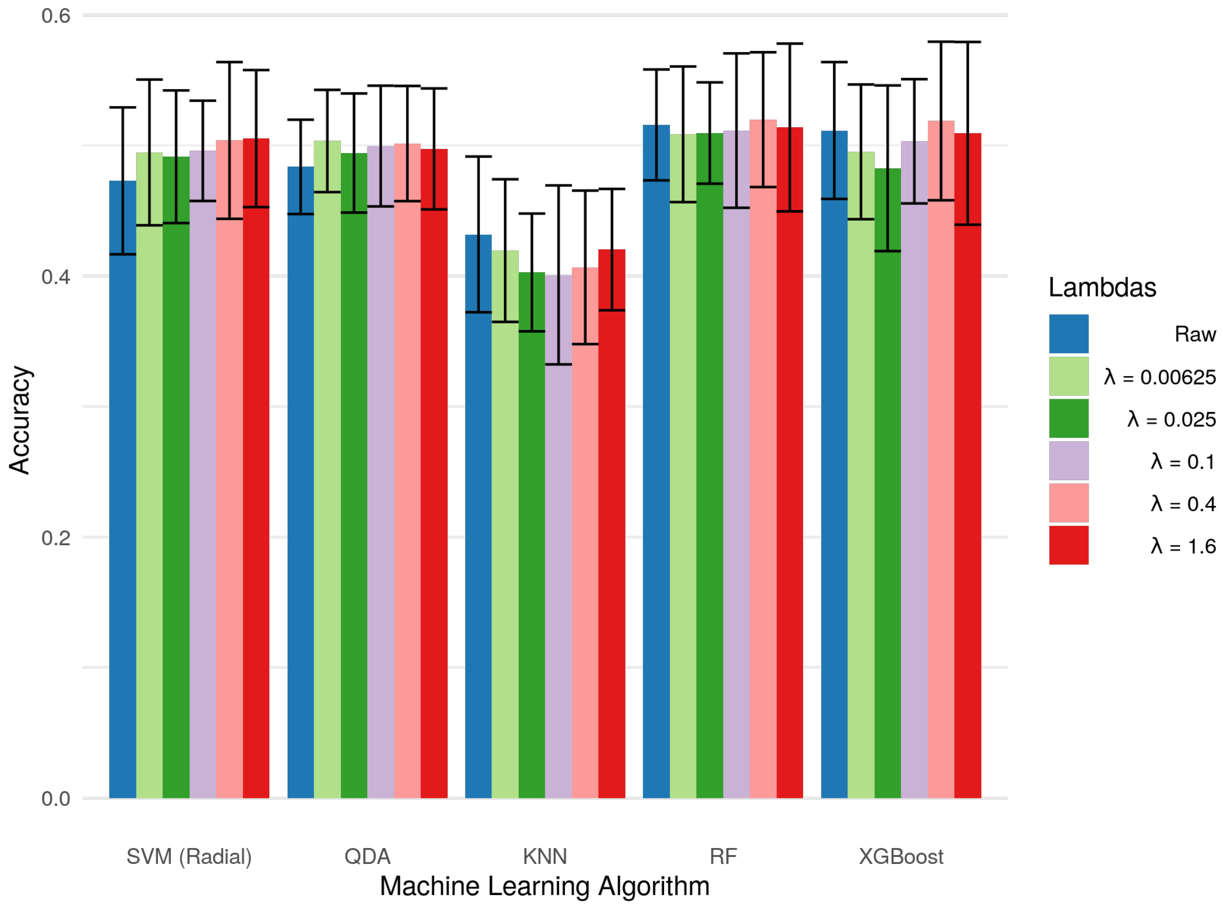


*Supplementary Figure 8: Bar graphs showing average accuracy for predicting Alzheimer disease status of various classifiers on raw and harmonized data across different hyperparameter choices. Error bars represent the standard deviation of validation-set accuracies across 5 repeats of 10-fold cross-validation.*

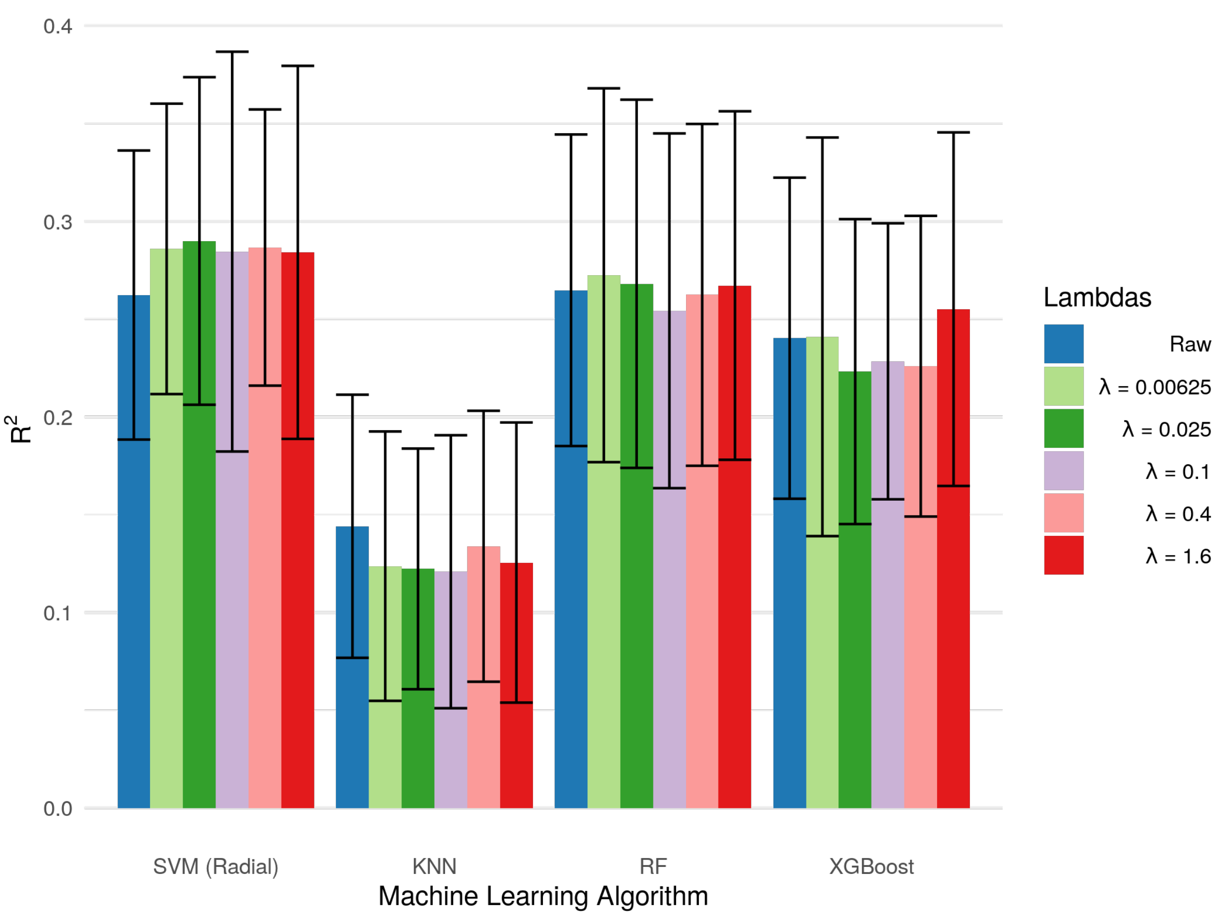


*Supplementary Figure 9: Bar graphs showing average R-squared value for predicting age of various classifiers on raw and harmonized data across different hyperparameter choices. Error bars represent the standard deviation of validation-set R-squared values across 5 repeats of 10-fold cross-validation.*
